# Multiomic deep delve of synthesis and secretion processes in a model peptidergic system

**DOI:** 10.1101/2022.05.31.494122

**Authors:** Soledad Bárez-López, André S. Mecawi, Natasha Bryan, Benjamin T. Gillard, Audrys G. Pauza, David Murphy, Michael P. Greenwood

## Abstract

The cell bodies of hypothalamic magnocellular neurones are densely packed in the hypothalamic supraoptic nucleus (SON) whereas their axons project to the anatomically discrete posterior pituitary gland. We have taken advantage of this unique anatomical structure to establish proteome and phosphoproteome dynamics in neuronal cell bodies and axonal terminals in response to physiological stimulation. We have found that proteome and phosphoproteome responses are very different between somatic and axonal neuronal compartments, indicating the need of each cell domain to differentially adapt. In particular, changes in the phosphoproteome in the cell body are involved in the reorganisation of the cytoskeleton and in axonal terminals the regulation of synaptic and secretory processes. We have identified that prohormone precursors including vasopressin and oxytocin are phosphorylated in axonal terminals and become hyperphosphorylated following stimulation. By multi-omic integration of transcriptome and proteomic data we identify changes to proteins present in afferent inputs to this nucleus.

**Graphical abstract:** 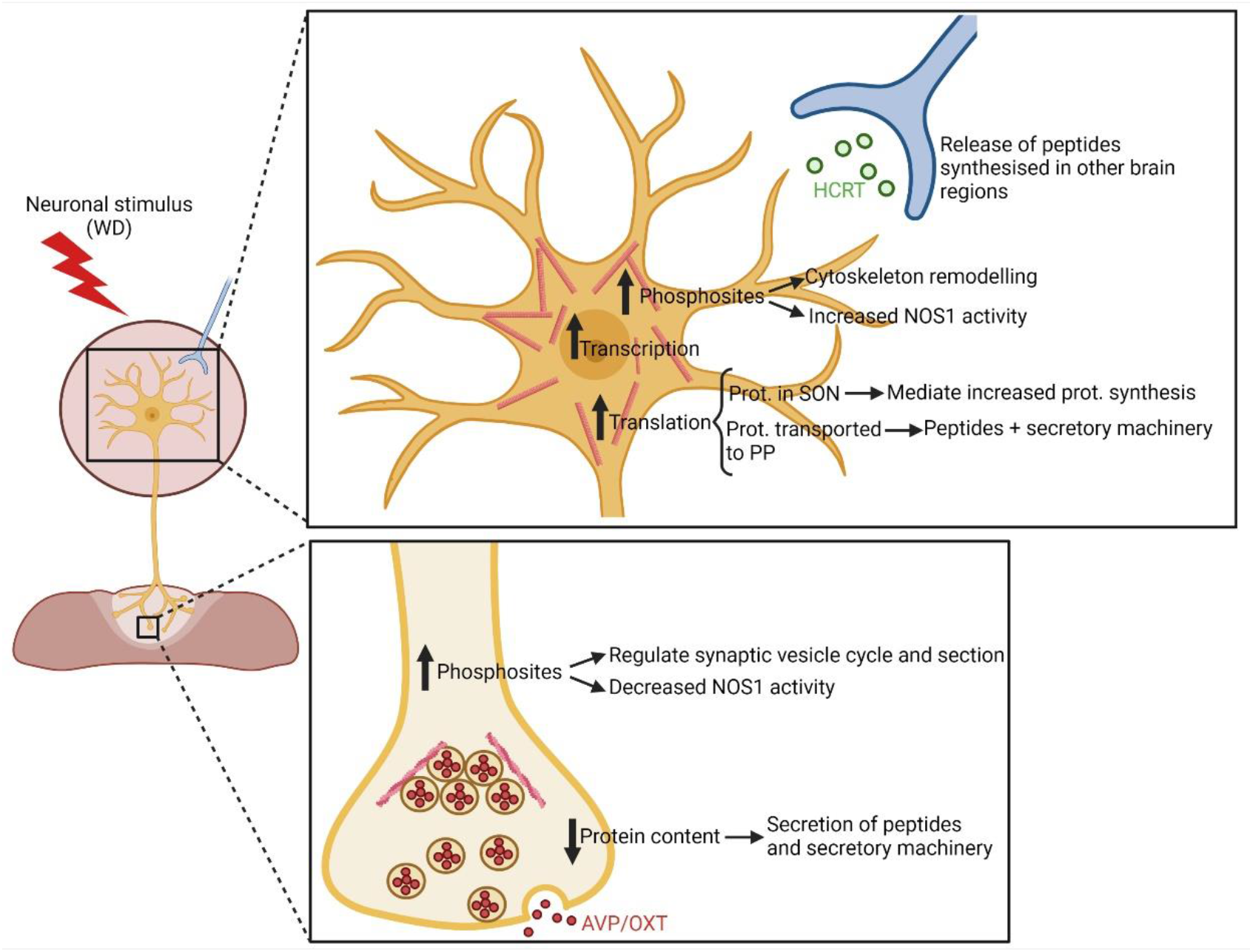

## Introduction

Determining the integrative transcriptome, proteome and phosphoproteome of a given cell type at any given time is rather challenging, but this is particularly complicated in neurones. It is recognised that neurones have distinct protein populations in their cell bodies and axons (Costa and Willis, 2018; Sahoo et al., 2018). Since proteins and their phosphorylation events control cellular function, it is imperative to separately determine the proteomes and phosphoproteomes of cell bodies and axonal terminals to better understand cellular processes in health and disease. The complex projection patterns of neurones makes this challenging for most brain nuclei.

Because of its unique anatomical structure, there is one neuronal system that is ideally suited to overcome many of these inherent challenges. The hypothalamo-neurohypophysial system (HNS) is a key neuroendocrine interface that connects the hormonal, neuronal, and vascular systems in all vertebrate species (Burbach et al., 2001; Mecawi et al., 2015). Hormones, such as antidiuretic hormone arginine vasopressin (AVP) and oxytocin (OXT), are made by densely packed populations of magnocellular neurones (MCN) predominantly located in the hypothalamic supraoptic (SON) and also the paraventricular nucleus (PVN). MCNs have one long axon that terminates in the posterior lobe of the pituitary gland (PP) and collaterals that project into other regions of the brain (Zhang et al., 2021). The PP is composed of pituicytes, approximately 30% by volume, and up to 36 million nerve terminals and swellings based on estimates of 18000 MCNs each with one long axon giving rise to an estimated 2000 nerve terminals and swellings (Leng and Ludwig, 2008). Each axon terminal contains ∼260 dense core vesicles packed with peptide hormones, including AVP and OXT, that are destined for release into the blood (Nordmann, 1977).

When the HNS is osmotically stimulated, such as evoked by water deprivation (WD), there is an increase in hormone release from the PP into the blood as the nerve endings are depolarised (Brownstein et al., 1980). Chronic osmotic stimulation of MCNs results in a striking functional remodelling of the HNS through several activity-dependent changes in the morphology, electrical properties and biosynthetic and secretory activity of the SON (Hatton, 1997; Sharman et al., 2004; Theodosis et al., 1998), which contribute to the facilitation of hormone synthesis, vesicle transportation, and secretion. Whilst this plasticity has been explored extensively at the transcriptome level (Dutra et al., 2021; Greenwood et al., 2015b; Hindmarch et al., 2006; Johnson et al., 2015; Pauža et al., 2021; Qiu et al., 2011), dynamic changes in the proteome and phosphoproteome have not thus far been addressed.

In this work we explore the proteomes and phosphoproteomes of the SON and the neurointermediate lobe (NIL) of the pituitary gland under basal and WD conditions. By integrating transcriptome catalogues, we comprehensively describe dynamic biosynthetic and secretory strategies that fuel the outputs of MCNs.

## Results

### Quantitative proteome and phosphoproteome of the rat SON

To stimulate MCNs, animals were WD for 48 hours. Stimulated animals were compared to euhydrated controls. The SON (containing MCN cell bodies and dendrites) was punched from the hypothalamus and the NIL (containing axonal terminals in the PP and the intermediate lobe) was dissected from the anterior lobe of the pituitary. Proteins were extracted and processed for Nano-LC Mass Spectrometry (LC-MS/MS, **Figure 1A**). A catalogue of proteins detected in the SON and NIL and differentially produced proteins and phosphosites between control and WD rats is presented in the supporting information (Tables S1 and S2).

**Figure 1.**
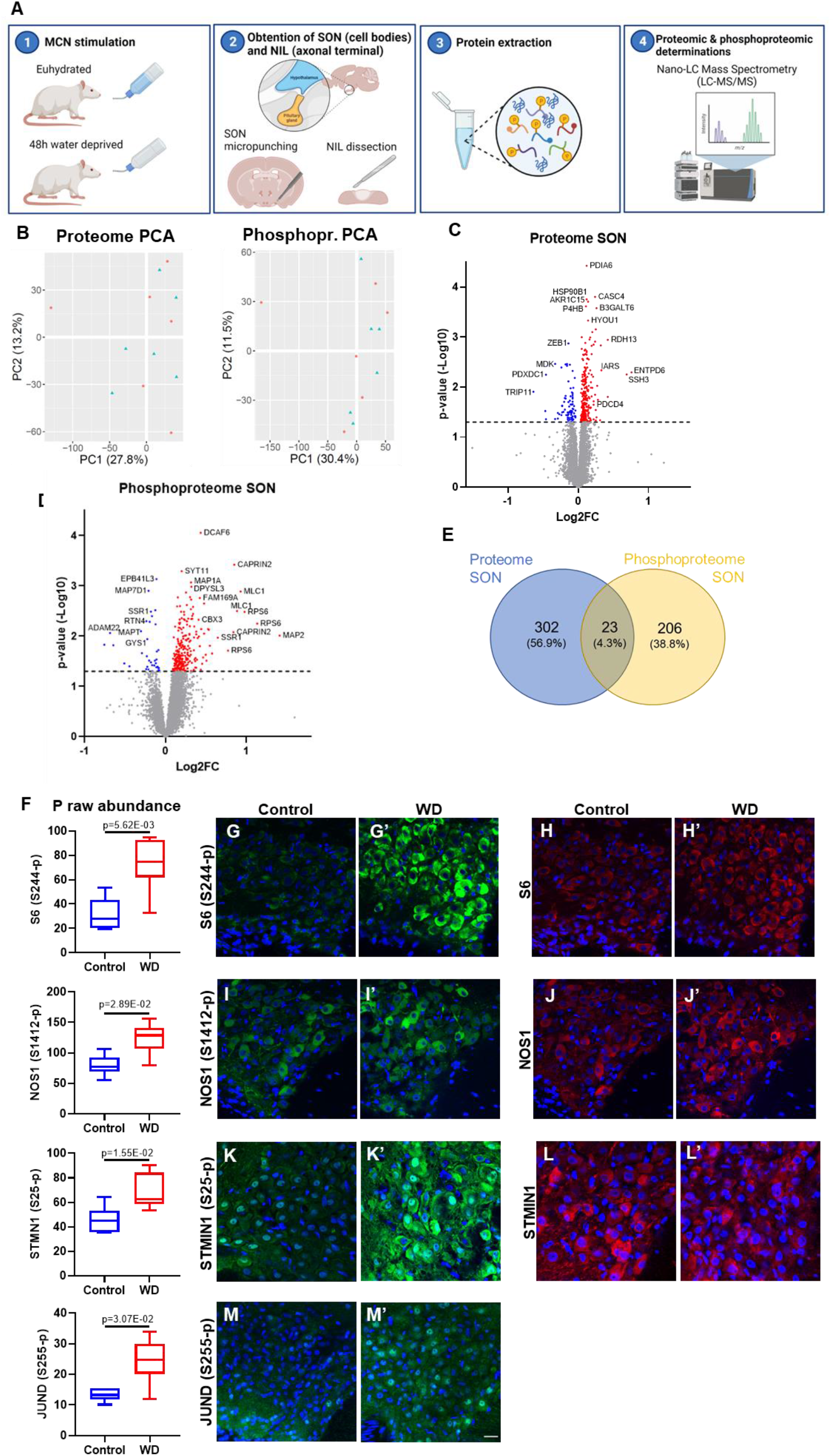
Quantitative proteome and phosphoproteome of the rat supraoptic nucleus (SON). (A) Graphical representation of the experimental approach. (1) 12 adult Sprague-Dawley rats were divided into 2 groups: 6 control with constant access to water and 6 subjected to a 48-hour water deprivation protocol (WD) to activate magnocellular neurones (MCNs). (2) The supraoptic nucleus (SON, mainly containing MCN cell bodies and dendrites) was punched from the hypothalamus and the neurointermediate lobe (NIL, mainly containing axonal terminals from the posterior pituitary as well as the intermediate lobe) was dissected from the anterior pituitary. (3) Proteins from SON and NIL were extracted and processed for Nano-LC Mass Spectrometry (LC-MS/MS). (4) Proteomics and phosphoproteomic determinations were performed by LC-MS/MS. Generated using BioRender (https://biorender.com/). (B) Principal component analysis (PCA) of the SON proteome and phosphoproteome in control (blue, n = 6) and WD rats (red; n = 6). (C) Volcano plot of WD vs control SON proteome showing 247 upregulated (p-value <0.05, red) and 78 downregulated (p-value <0.05, blue) proteins. (D) Volcano plot of WD vs control SON phosphoproteome showing 252 hyperphosphorylation (p-value <0.05, red) and 36 hypophosphorylation (p-value <0.05, blue) events. (E) Venn diagram showing 23 proteins in common with changes at the proteome and phosphoproteome level in response to WD. (F) Phospho raw abundance for S244-p S6, S1412-p NOS1, S25-p STMN1 and S255-p JUND in the SON according to LC-MS/MS between control (n = 6) and WD (n = 6) rats. (G) Immunohistochemistry against S244-p S6 in the SON of control and (G’) WD rats. (H) Immunohistochemistry against S6 in the SON of control and (H’) WD rats. (I) Immunohistochemistry against S1412-p NOS1 in the SON of control and (I’) WD rats. (J) Immunohistochemistry against NOS1 in the SON of control and (J’) WD rats. (K) Immunohistochemistry against S25-p STMN1 in the SON of control and (K’) WD rats. (L) Immunohistochemistry against STMN1 in the SON of control and (L’) WD rats. (M) Immunohistochemistry against S255-p JUND in the SON of control and (M’) WD rats. Images are representative of n = 4. Scale bar represents 25 µm.

Proteome analysis of the SON identified 7668 proteins (Table S1). Principal component analysis (PCA) of proteome and phosphoproteome data did not reveal a separation pattern between control and WD groups (**Figure 1B**). Of the 7668 proteins detected in the SON, the data indicated 325 differentially abundant proteins in WD SON, with 247 being increased and 78 decreased in content (**Figure 1C**). We asked about the phosphorylation status of identified proteins. We found that 229 proteins underwent changes in phosphorylation in response to WD (p-value < 0.05), which included 252 hyperphosphorylation and 36 hypophosphorylation events (**Figure 1D**). Only 23 proteins that changed in overall content also had phosphorylation modifications in response to WD (**Figure 1E**).

The phosphoproteome data was validated using commercially available phopho-antibodies against significantly altered phosphosites in the control and WD SON. We investigated phosphosites for 40S ribosomal protein S6 (S6; S244-p) that showed a Log2 Fold Change (Log2FC) of 1.13 (p-value = 5.62E-03), Nitric oxide synthase (NOS1, S1412-p) with a Log2FC of 0.36 (p-value = 2.89E-02), STATHMIN1 (STMN1, S25-p) with a Log2FC of 0.55 (p-value = 1.55E-02) and JUND (S255-p) with a Log2FC of 0.51 (p-value = 3.07E-02) (**Figure 1F**). In addition, we immunolabelled S6, NOS1 and STMN1 proteins. Immunohistochemical studies supported increased phosphorylation of S6 S244-p in MCNs (**Figure 1G, G’**) with no change to S6 protein content (**Figure 1H, H’**). The phosphorylation of the S1412-p residue of NOS1 (**Figure 1I, I’**) appeared to increase in a population of MCNs that produce NOS1 (**Figure 1J, J’**). Immunostaining against S25-p STMN1, detected the presence of this phospho residue in the SON and revealed increased staining in MCNs in response to WD (**Figure 1K, K’**). No differences were observed in immunostaining of the STMN1 protein (**Figure 1L, L’**). Immunostaining of S255-p JUND revealed the presence of this phospho residue in the SON and confirmed an increase in phosphorylation in MCNs in WD (**Figure 1M, M’**). We have previously reported increased JUND protein content by western blot and immunohistochemistry in the WD SON (Yao et al., 2012). All these findings agreed with the LC-MS/MS output.

### Quantitative proteome and phosphoproteome of the rat NIL

Proteome analysis identified 9303 proteins in the NIL (Table S2). PCA using all proteins revealed distinct separation between the total proteomes of control and WD samples with PC1 accounting for 26% of total variance between samples, which was attributable to WD (**Figure 2A**). PCA also displayed a distinctive separation between the phosphoproteomes of control and WD samples with PC1 explaining 19.8% of the total variance between samples, that was attributable to the experimental condition (**Figure 2A**). Out of 9303 total proteins, 870 changed their protein content as a consequence of WD (p-value < 0.05) with 282 proteins decreasing their total content and 588 increasing their total protein content (**Figure 2B**). We again asked about the phosphorylation status of identified proteins. We found that 760 proteins underwent changes in phosphorylation (p-value < 0.05), which included 746 hyperphosphorylation and 755 hypophosphorylation events (**Figure 2C**). A total of 151 proteins with alterations to their overall content also underwent changes in phosphorylation in WD (**Figure 2D**).

**Figure 2.**
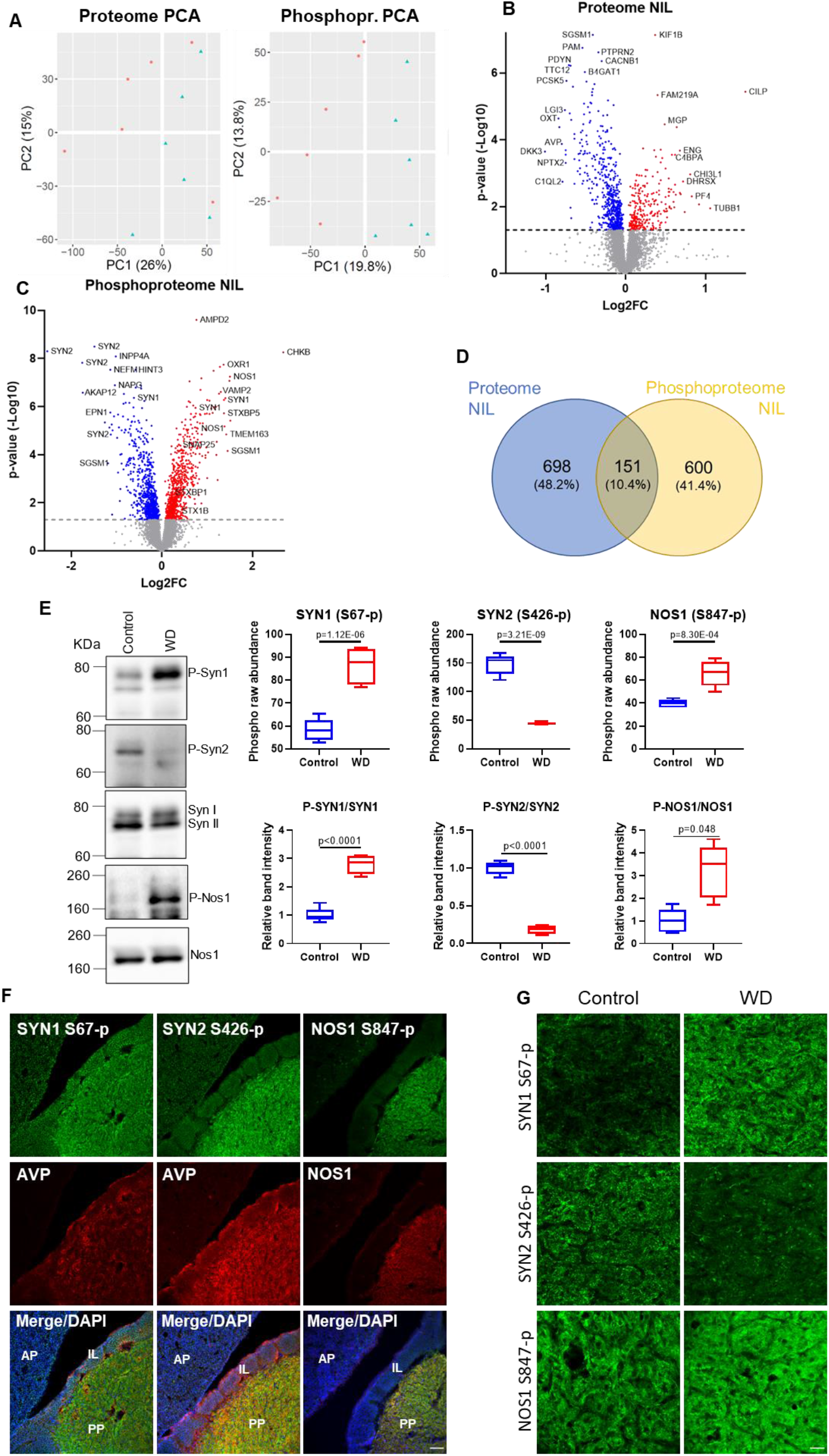
Quantitative proteome and phosphoproteome of the rat neurointermediate lobe (NIL) (A) Principal component analysis (PCA) of the NIL proteome and phosphoproteome in control (blue, n = 6) and water-deprived (WD) rats (red; n = 6). (B) Volcano plot of WD vs control NIL proteome showing 276 upregulated (p-value <0.05, red) and 573 downregulated (p-value <0.05, blue) proteins. (C) Volcano plot of WD vs control NIL phosphoproteome showing 746 hyperphosphorylation (p-value <0.05, red) and 755 hypophosphorylation (p-value <0.05, blue) events. (D) Venn diagram showing 151 proteins in common with changes at the proteome and phosphoproteome level in response to WD (E) Phospho raw abundance for S67-p SYN1, S426-p SYN2 and S426-p SYN2 in the NIL according to LC-MS/MS between control (n = 6) and WD (n = 6) rats. Western blotting analysis of S67-p SYN1 (normalised against SYN1), S426-p SYN2 (normalised against SYN2) and S847-p NOS1 (normalised against NOS1) in control (n = 5) and WD NILs (n = 5 for S847-p NOS1 and n = 4 for S67-p SYN1 and S426-p SYN2). (F) Immunohistochemistry against S67-p SYN1 and arginine vasopressin (AVP), S426-p SYN2 and AVP and S847-p NOS1 and NOS1 in the pituitary gland of control rats showing the anterior pituitary (AP), intermediate lobe (IL) and posterior pituitary (PP). Images are representative of n = 4. Scale bar represents 75 µm. (G) Immunohistochemistry against S67-p SYN1, S426-p SYN2 and S847-p NOS1 in the PP of control and WD rats. Images are representative of n = 4. Scale bar represents 25 µm.

NIL phosphoproteome data was validated by western blot and immunohistochemistry. We used commercially available phospho-antibodies against the phosphosites of Synapsin 1 (SYN1, S67-p) that showed a Log2FC of 0.74 (p-value = 1.12E-06), Synapsin 2 (SYN2, S426-p) with a Log2FC of -1.49 (p-value = 3.21E-09), and NOS1 (S847-p) with a Log2FC of 0.67 (p-value = 8.30E-04) (**Figure 2E**), as well as antibodies against SYN and NOS1 proteins. Immunoblotting confirmed an increase in SYN1 S67-p, decrease in SYN2 S426-p, and increased NOS1 S847-p following WD (**Figure 2E; S1)**. Immunostaining showed the location of these phosphosites predominantly in the PP which was marked with AVP (**Figure 2F**). Higher magnification images of the PP in WD confirmed an increase in SYN1 S67-p, a decrease in SYN2 S426-p, and an increase in NOS1 S847-p **Figure 2G**). Overall, the pattern of phosphoregulation observed in the SON and NIL agreed with the LC-MS/MS output.

### Pathway analyses and functional classification of the proteomes and phosphoproteomes of SON and NIL

To gain functional insights into the responses of the MCN compartments localised in the SON and NIL to WD, we performed pathway analysis of the differential produced proteins between control and WD samples by interrogating GO and KEGG databases. We show enriched terms (restricted to up to 15 terms) retrieved for each category ranked according to P_Adj_ value (**Figure 3**). In addition, we plotted the topmost significant associated differentially produced proteins coloured based on Log2FC and sized according to total normalised protein following WD. In the SON, all the terms retrieved for the GO:Cellular Compartment Process (GO:CC) and GO:Biological Process (GO:BP) categories highlighted terms associated with the endomembrane system including the endoplasmic reticulum (ER), the Golgi apparatus, as well as Golgi-associated vesicles and COPI-coated vesicles (**Figure 3A** and **Table S3**). No enriched terms were identified in the GO:Molecular Function Process (GO:MF) category. Over-representation analysis by KEGG identified “Protein processing in endoplasmic reticulum” as an enriched pathway (**Figure 3A** and **Table S3**). Strikingly, most of the top-regulated proteins increased their content in response to WD. Amongst the most abundant proteins following WD were a series of chaperones known to regulate protein folding and stress in the ER including the Heat Shock Protein Family A (Hsp70) Member 5 (HSPA5) also known as Endoplasmic reticulum chaperone BiP, the Hypoxia up-regulated protein 1 (HYOU1), Protein disulfide-isomerase (P4HB) and calreticulin (CALR) (Ni and Lee, 2007).

**Figure 3.**
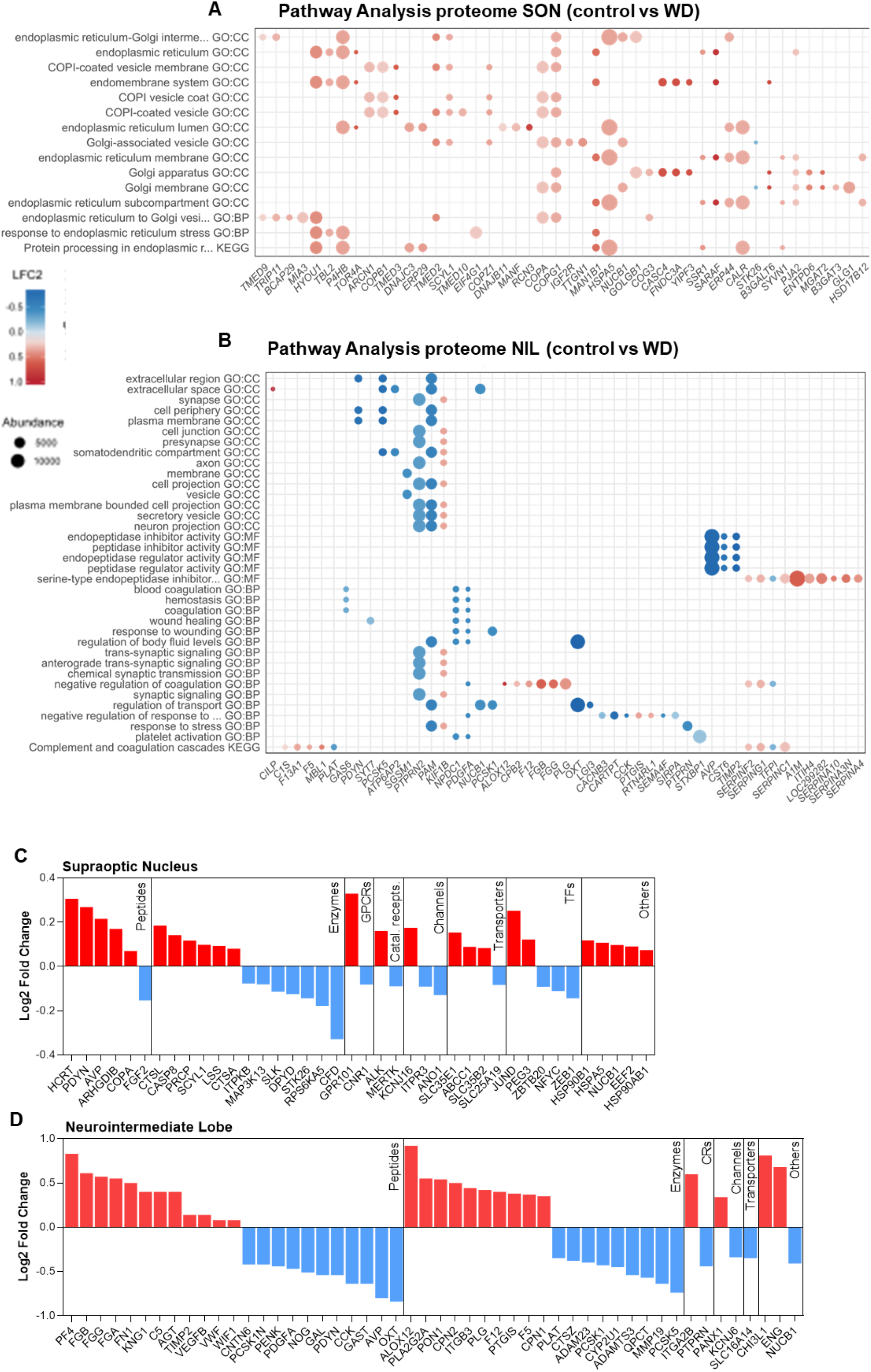
Pathway analyses and functional classification of the proteomes of supraoptic nucleus (SON) and neurointermediate lobe (NIL) (A) Pathway analysis of changes in the SON proteome as a result of water deprivation (WD) using GO and KEGG databases. Dot plot of all enriched terms retrieved for each category ranked according to P_Adj_ value from top to bottom in increasing order. The top 10 most significant associated differentially expressed proteins of each over-represented category are shown as dots coloured based on Log2FC and sized according to total normalised protein expression following WD. (B) Pathway analysis of changes in the NIL proteome as a result of WD using GO and KEGG databases. Dot plot of up to 15 enriched terms retrieved for each category ranked according to P_Adj_ value from top to bottom in increasing order. The top 10 most significant associated differentially expressed proteins of each over-represented category are shown as dots coloured based on Log2FC and sized according to total normalised protein expression following WD. (C) Proteome Log2FC changes in the rat SON as a consequence of WD categorised according to their pharmacological classification or their function as a transcription factor. (D) Proteome Log2FC changes in the rat NIL as a consequence of WD categorised according to their pharmacological classification or their function as a transcription factor.

In the NIL, terms highlighted in the GO:CC category included “synapse”, “presynapse” and “secretory vesicle”, amongst others (**Figure 3B** and **Table S3**). In the GO:MF hierarchy and the KEGG pathway, all terms were related to endopeptidase activity, probably triggered by the changes in hormone content, shown here by AVP, in response to WD (**Figure 3B** and **Table S3**). The GO:BP included several terms related to wounding and coagulation, but more pertinent to the present study were the several terms related to synaptic signalling, regulation of transport or regulation of body fluid levels (**Figure 3B** and **Table S3**). KEGG analysis identified the enriched pathway “Complement and coagulation cascades” (**Figure 3B** and **Table S3**). The coagulation-related terms identified in the GO:BP and KEGG analysis, such as coagulation factor V (F5), Platelet-derived growth factor subunit A (PDGFA) or Plasma protease C1 inhibitor (SERPING1), could be related to dehydration-mediated blood coagulation activation (Shi et al., 2019) or to the increase in perivascular protrusions that have been described in the PP following chronic stimulation (Miyata, 2017). Some of the proteins with biggest Log2FC included those involved in “regulation of fluid levels”, such as the peptides AVP and OXT or the enzyme peptidylglycine alpha-amidating monooxygenase (PAM), that mediates C-amidation of endogenous peptides including AVP and OXT (Yin et al., 2011). Other proteins with large Log2FC were serine-protein kinase ATM (ATM) involved in vesicle transport (Pizzamiglio et al., 2020) or receptor-type tyrosine-protein phosphatase N2 (PTPRN2), also involved in vesicle-mediated secretory processes (Wasmeier et al., 2005). Altogether, these data suggests that changes in the proteome in response to WD in the SON are associated with protein synthesis, whilst in the NIL they are related to synaptic signalling and transport.

To further investigate differentially produced proteins in separate compartments of the HNS, we mined the IUPHAR (Harding et al., 2022) and a transcription factor database (Lambert et al., 2018) to report physiological and pharmacological classifications or identity as transcription factors. In the SON (**Figure 3C**), 6 differentially synthesised proteins were classified as peptides, 5 increasing and 1 decreasing in protein content. Notably, this included increased AVP and Proenkephalin-B (PDYN) which are known to be increased in MCNs of the WD SON (Pauža *et al*., 2021). The largest category was enzymes which was much more evenly weighted with proteins increasing and decreasing in content. In the NIL (**Figure 3D**), a series of peptides increased their content while others, such as OXT, AVP, and PDYN decreased, illustrating the known role for the PP in peptide secretion (Brown, 2016). Several enzymes also changed their content in the NIL in response to WD, but none of these coincided with those in the SON, suggesting different adaptation to WD between these two structures.

We next performed pathway analysis using GO and KEGG databases to explore the phosphoproteome changes as a result of WD in the SON and NIL. We ranked according to P_Adj_ value up to 15 of the enriched terms identified in each category and we plotted the proteins with most significant phosphorylation changes in a phosphosite. In addition, we indicate the total number of phosphorylation events in that protein following WD (**Figure 4**). In the SON, the only term retrieved in the GO:CC category was “cell junction” whilst both the GO:MF and the GO:BP hierarchies highlighted terms related to microtubule and cytoskeleton binding and organisation (**Figure 4A** and **Table S3**). We identified phosphorylation changes in several microtubule-associated proteins (MAPs), organisers of the microtubule cytoskeleton (Bodakuntla et al., 2019), including MAP2, MAP1A and MAPT. We also found phosphorylation events in STMN1, known to control microtubule dynamics (Benarroch, 2021; Rubin and Atweh, 2004) and other microtubule-organising proteins such as Calmodulin-regulated spectrin-associated protein 2 (CAMSAP2) (Yau et al., 2014) and Microtubule crosslinking factor 1 (MTCL1). No significantly enriched terms were identified in the KEGG pathways.

**Figure 4.**
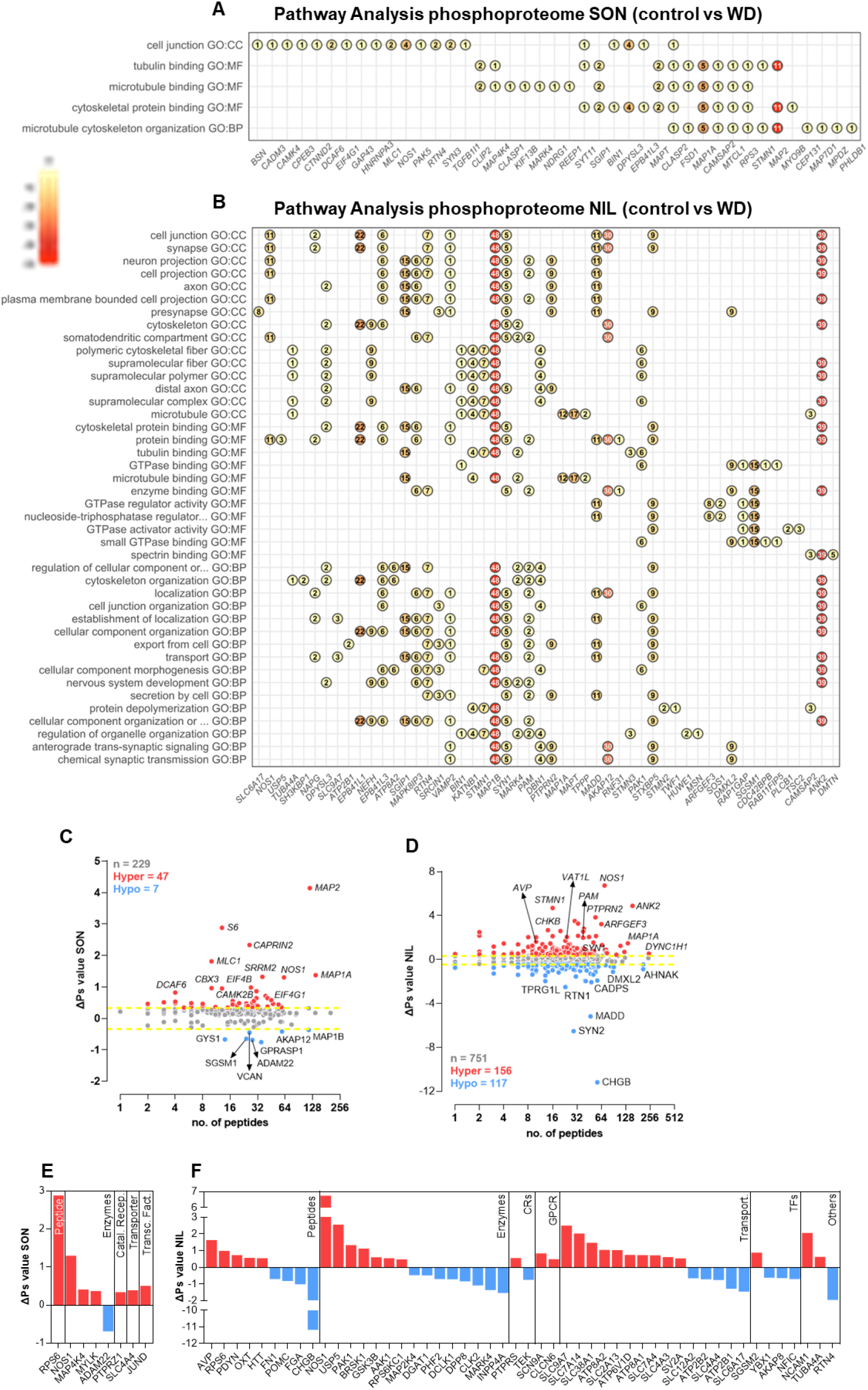
Pathway analyses and functional classification of the phosphoproteomes of supraoptic nucleus (SON) and neurointermediate lobe (NIL) (A) Pathway analysis of changes in the SON phosphoproteome as a result of water deprivation (WD) using GO and KEGG databases. Dot plot of all enriched terms retrieved for each category ranked according to P_Adj_ value from top to bottom in increasing order. The top 20 proteins with most significant phosphorylation changes in a phosphosite are shown as a dot indicating the total number of phosphorylation events in that protein following WD. (B) Pathway analysis of changes in the NIL phosphoproteome as a result of WD using GO and KEGG databases. Dot plot of up to 15 enriched terms retrieved for each category ranked according to P_Adj_ value from top to bottom in increasing order. The top 15 proteins with most significant phosphorylation changes in a phosphosite are shown as a dot indicating the total number of phosphorylation events in that protein following WD. (C) Global overall phosphorylation state change (ΔPs) analysis of phosphoproteins between control and WD rats in the SON. Numbers of hyperphosphorylated (Hyper) and hypophosphorylated (Hypo) peptides are shown. Dotted lines, ΔPs = ±0.34. (D) Global ΔPs analysis of phosphoproteins between control and WD rats in the NIL. Numbers of hyperphosphorylated (Hyper) and hypophosphorylated (Hypo) peptides are shown. Dotted lines, ΔPs = ±0.4. (E) ΔPs changes in the rat SON as a consequence of WD categorised according to their pharmacological classification or their function as a transcription factor. (F) ΔPs changes in the rat NIL as a consequence of WD categorised according to their pharmacological classification or their function as a transcription factor.

In the NIL the GO:CC highlighted several terms including “synapse”, “presynapse” and “cytoskeleton”. The GO:MF category retrieved several terms related to protein binding and the GO:BP hierarchy highlighted terms related to cellular organisation and localisation in addition to other terms related to secretion and synaptic signalling and transmission (**Figure 4B** and **Table S3**). As such, phosphorylated proteins were mainly involved in cytoskeleton organisation and localisation of cellular components including MAP1B, MAP1A, Ankyrin-2 (ANK2) and Band 4.1-like protein 1 (EPB41L1) or in secretion and synaptic transmission including vesicle-associated membrane protein 2 (VAMP2), SYN1 and Syntaxin-binding protein 5 (STXBP5). No enriched terms were identified in the KEGG pathways. We thus establish that changes in phosphorylation in response to WD in the SON are involved in cytoskeleton organisation whereas in the NIL they also regulate synaptic and secretory processes.

We then globally quantified changes in phosphorylation in order to obtain single values for changes in the overall phosphorylation status of individual proteins. This analysis measured the overall phosphorylation state change (ΔPs) of each protein as described by Wang et al. (2018). In the SON, the ΔPs showed that 47 proteins were significantly hyperphosphorylated (ΔPs > 0.34) whilst 7 were hypophosphorylated (ΔPs < -0.34) (**Figure 4C**). The top 3 ΔPs hyperphosphorylated proteins were MAP2, S6 and Cytoplasmic activation/proliferation-associated protein 2 (CAPRIN2) and the hypophosphorylated proteins were G-protein coupled receptor-associated sorting protein 1 (GPRASP1), Disintegrin and metalloproteinase domain-containing protein 22 (ADAM22) and Glycogen synthase (GYS1). In the NIL, the ΔPs analysis revealed 157 significant hyperphosphorylated (ΔPs > 0.4) proteins and 117 hypophosphorylated proteins (ΔPs < 0.4) (**Figure 4D**). The top 3 ΔPs hyperphosphorylated proteins were NOS1, STMN1 and ANK2 while the hypophosphorylated ones included Secretogranin-1 (CHGB), SYN2 and MAP kinase-activating death domain protein (MADD).

We then plotted the proteins with significant changes in ΔPs in response to WD and classified them according to their physiological and pharmacological classifications or their identity as transcription factors in the SON (**Figure 4E**) and the NIL (**Figure 4F**). The ΔPs changed in only 1 peptide in the SON whilst several peptides underwent changes in phosphorylation in the NIL, including AVP, OXT and PDYN, all of which decreased their total protein content in the NIL following WD (**Figure 4D**). This finding strongly suggests that changes in peptide phosphorylation in the NIL might be related to peptide secretion. Several enzymes had changed ΔPs in the SON and NIL including, of note, NOS1 which was hyperphosphorylated in both structures (**Figure 4C, D** and **S2**). Additionally, various transporters had significant changes in their ΔPs in the NIL, with only 1 in the SON.

### Phosphorylation adaptations in the SON and NIL in response to WD

Pathway analysis of the changes in the phosphoproteome following WD were indicative of important differential adaptations in the SON and the NIL. Phosphorylation modifications in the SON seemed to mediate cytoskeleton remodelling whilst in the NIL they were related to synaptic events. To further explore post-translational events elicited by WD, we mapped the phosphorylation sites identified by LC-MS/MS to proteins involved in selected ontology terms.

In the SON, we explored the only GO:BP term retrieved: “microtubule cytoskeleton organisation” (**Figure 4A**). We mapped all phosphosites detected in this structure by LC-MS/MS highlighting those that underwent hyper or hypophosphorylation events in selected proteins from this GO category (**Figure 5**). These included Centrosomal protein of 131 kDa (CEP131), Mapb1, Cytoplasmic Linker Associated Protein 2 (CLASP2), Multiple PDZ domain protein (MPDZ), Pleckstrin homology-like domain family B member 1 (PHLDB1), CAMSAP2, MAP2, MTCL1, STMN1, MAP1A, MAP7 domain-containing protein 1 (MAP7D1) and Microtubule Affinity Regulating Kinase 4 (MARK4), among others. All these proteins were hyperphosphorylated, with the exception of MAP7D1 that was hypophosphylated and MAP1B that was both hyper and hypophosphorylated at different residues. To further explore the phosphorylation events in synapse-related categories in the NIL, we used SynGO analysis (Koopmans et al., 2019) to detail synapse-specific changes. For the cellular compartment ontology, the most enriched terms revealed were at the level of the presynapse (**Figure 6A**). Biological Process ontology terms highlighted terms such as “process in the presynapse”, “synaptic vesicle cycle” and “presynaptic dense core vesicle (DCV) exocytosis” (**Figure 6B, Table S4**). We mapped all phosphosites detected for some of the proteins involved in the “synaptic vesicle cycle” category, which can be further classified into terms “regulation of synaptic vesicle cycle” composed of the proteins Bassoon (BSN), SYN1 and Rho GDP Dissociation Inhibitor Alpha (ARHGDIA), “synaptic vesicle clustering” comprising Piccolo (PCLO), Synapsin 3 (SYN3), SYN2, Abelson interactor 1 (ABI1) and BSN and “synaptic vesicle exocytosis” which included the proteins Synaptosomal-Associated Protein, 25kDa (SNAP25), VAMP2, unc-13 homolog A (UNC13A), Syntaxin 1B (STX1B) and Synaptotagmin 2 (SYT2), amongst others (**Figure 6C**). For the term “presynaptic dense core vesicle exocytosis” we mapped the phosphosites for some of the proteins in this category which included Calcium-dependent secretion activator 1 (CADPS), Regulating synaptic membrane exocytosis protein 1 (RIMS1), SNAP25, Dynamin-1 (DNM1), Syntaxin-binding protein 1 (STXBP1), STXBP5 and Ras-related protein Rab-3A (RAB3A) (**Figure 6C**). By mapping the phosphosites to these proteins as well as the hyperphosphorylation and hypophosphorylation sites, we highlight a series of phosphorylation events potentially involved in cytoskeleton organisation, the synaptic vesicle cycle and DCV exocytosis.

**Figure 5.**
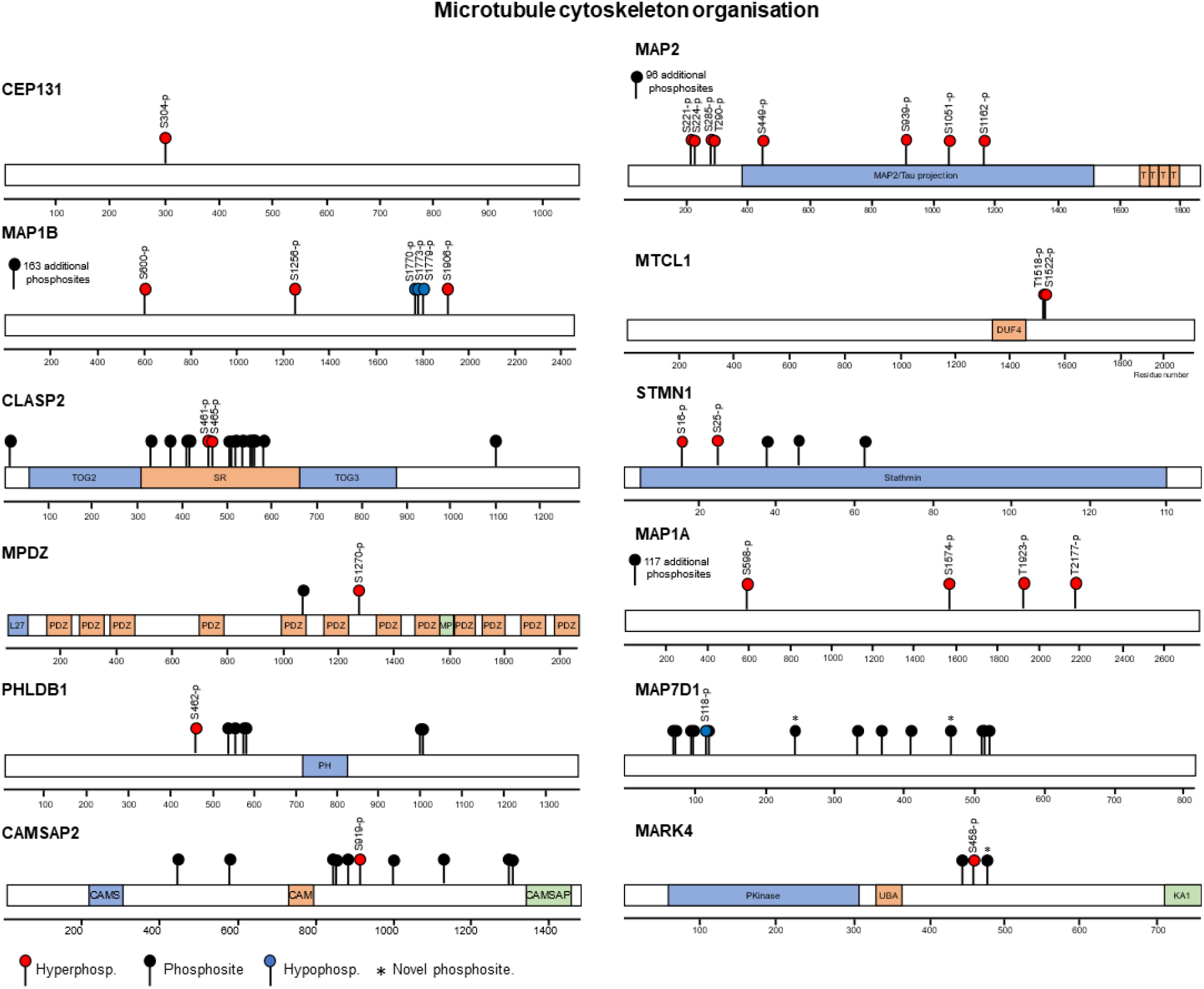
Phosphorylation events regulating cytoskeletal remodelling in the stimulated supraoptic nucleus. Mapping the phosphosites and hyper and hypophosphorylation events in response to water deprivation (WD) in proteins involved in microtubule cytoskeleton organisation. Protein domains include CA: Spectrin-binding region of Ca^2+^-Calmodulin, CAMS: CAMSAP Calponin-homology domain, CAMSAP: Calmodulin-regulated spectrin-associated CKK domain, DUF4: DUF4482 Domain of unknown function, KA1: Kinase associated domain 1, L27: L27 domain, MP: Unstructured region 10 on multiple PDZ protein, PDZ: post synaptic density protein (PSD95) domain, PH: Pleckstrin homology domain, PKinase: Protein kinase domain, SLD: Stathmin-like domain, SR: serine-arginine-rich region, T: Tau and MAP protein tubulin-binding repeat, TOG1,2: tumour overexpressed gene domains, UBA: ubiquitin-associated domain.

**Figure 6.**
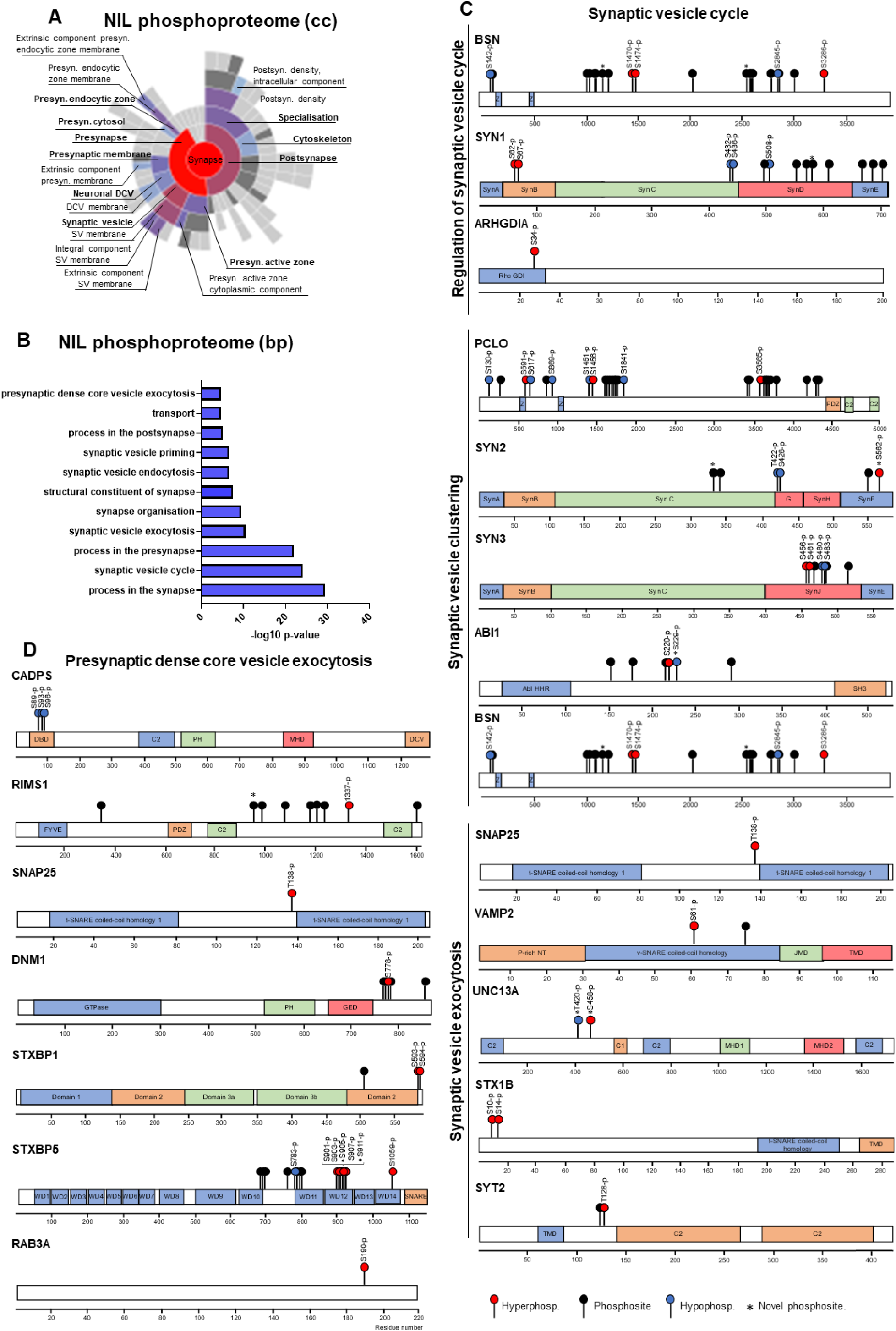
Phosphorylation events regulating synaptic processes in the stimulated neurointermediate lobe (NIL) (A) SynGO cellular compartment (cc) enrichment analysis of all the proteins undergoing phosphorylation events in response to water deprivation (WD) in the NIL. (B) SynGO biological processes (bp) enrichment analysis of all the proteins undergoing phosphorylation events in response to WD in the NIL. (C) Mapping the phosphosites and hyper and hypophosphorylation events in response to WD in proteins involved in the synaptic vesicle cycle including the regulation of the synaptic vesicle cycle, synaptic vesicle clustering and synaptic vesicle exocytosis and in proteins involved in presynaptic dense core vesicle exocytosis. Protein domains include AbI HHR: Abl-interactor homeo-domain homologous region, C1: phorbol esters/diacylglycerol binding domain C1 domain, C2: Ca^2+^-dependent C2 domain, DBD: dynactin 1 binding domain, DCV: dense core vesicle association domain, FYVE: FYVE zinc finger domain, GED: Dynamin GTPase effector domain, GTPase: GTPase domain, JMD: juxta-membrane domain, MHD1,2: Munc13-homology domains, P-rich NT: proline-rich N-terminal domain, PDZ: post synaptic density domain, PH: Pleckstrin homology domain, Rho GDI: RHO protein GDP dissociation inhibitor, Syn A, B, C, D, E, F, G, H, J: Synapsin domain A, B, C, D, E, F, G, H, J, TMD: transmembrane domain, WD: WD40 repeat or beta-transducin repeat, Z: Piccolo Zn-finger.

### Multiomic integration of stimulated SON and NIL

Next, we performed an integrative analysis of the differential transcriptomes, proteomes and phosphoproteomes of the SON and NIL by Spearman correlation analysis. This was done by comparing the proteome and phosphoproteome data with transcriptomic analysis of the 72-hour WD Wistar Han rats (Pauža et al., 2021). Spearman correlation analysis between differentially expressed genes and proteins in the SON in response to WD revealed a positive correlation (r = 0.55, p-value < 0.0001; **Figure 7A**) indicating that, in response to a stimulus such as WD, increased steady-state transcript abundance in general leads to increased translation in the SON. There was no such correlation between differentially expressed genes in the SON and differentially produced proteins in the NIL in response to WD (Spearman r = -0.099, p-value = 0.368; **Figure 7B**). However, there were a number of proteins with increased gene expression in the SON and decreased protein content in the NIL (such as AVP, PDYN, PCSK1 or PCSK5). This illustrates how the SON synthesises proteins which are then transported for release from the PP in response to stimulation. Moreover, the changes in total proteome between the SON and NIL in response to WD did not correlate (Spearman r = -0.209, p-value = 0.285; **Figure 7C**). A similar pattern was observed with the phosphoproteomes, where the ΔPs between the SON and the NIL did not show a statistically significant correlation (Spearman r = 0.140, p-value = 0.171; **Figure 7D**). This implies cell compartment specific changes in response to WD. We next explored the relationship between changes in ΔPs in the NIL and changes in the total proteome in response to WD in the SON. Spearman correlation analysis revealed a positive correlation (Spearman r = 0.495, p-value = 0.014; **Figure 7E**). This suggests that, following WD, the SON synthesises proteins that are transported to the NIL, where they are hyperphosphorylated. In addition, exploring the relationship between changes in ΔPs in the NIL in response to WD and the changes in the total proteome in response to WD in the NIL showed a negative correlation (Spearman r = -0.518, p-value = 0.0001; **Figure 7F**) suggesting that, in response to a stimulus such as WD, hyperphosphorylated proteins are secreted from the NIL into the circulation or are degraded. Among the proteins that are hyperphosphorylated and secreted from the NIL were the peptides AVP, OXT, PDYN and the enzyme PAM responsible for peptide C-amidation (Yin *et al*., 2011). To further explore the possible role of hyperphosphorylation in the secretion of these proteins, we mapped the identified phosphosites in the SON and the NIL to the protein sequence (**Figure 7G**). Interestingly, in the SON, none of these proteins underwent any changes in phosphorylation to WD, indeed they presented no phosphorylation events at all. In the NIL we detected 8 phosphosites (7 of them described in the present the work from the first time), 4 of which were hyperphosphorylated in response to WD. Two novel sites were found for OXT and one of them was hyperphosphorylated in response to WD. For PDYN we identified 6 phosphosites, 3 of which were hyperphophorylated following stimulus. Of the 2 phosphosites identified in PAM in the NIL, one was hyperphosphorylated as a consequence of neuronal activation.

**Figure 7.**
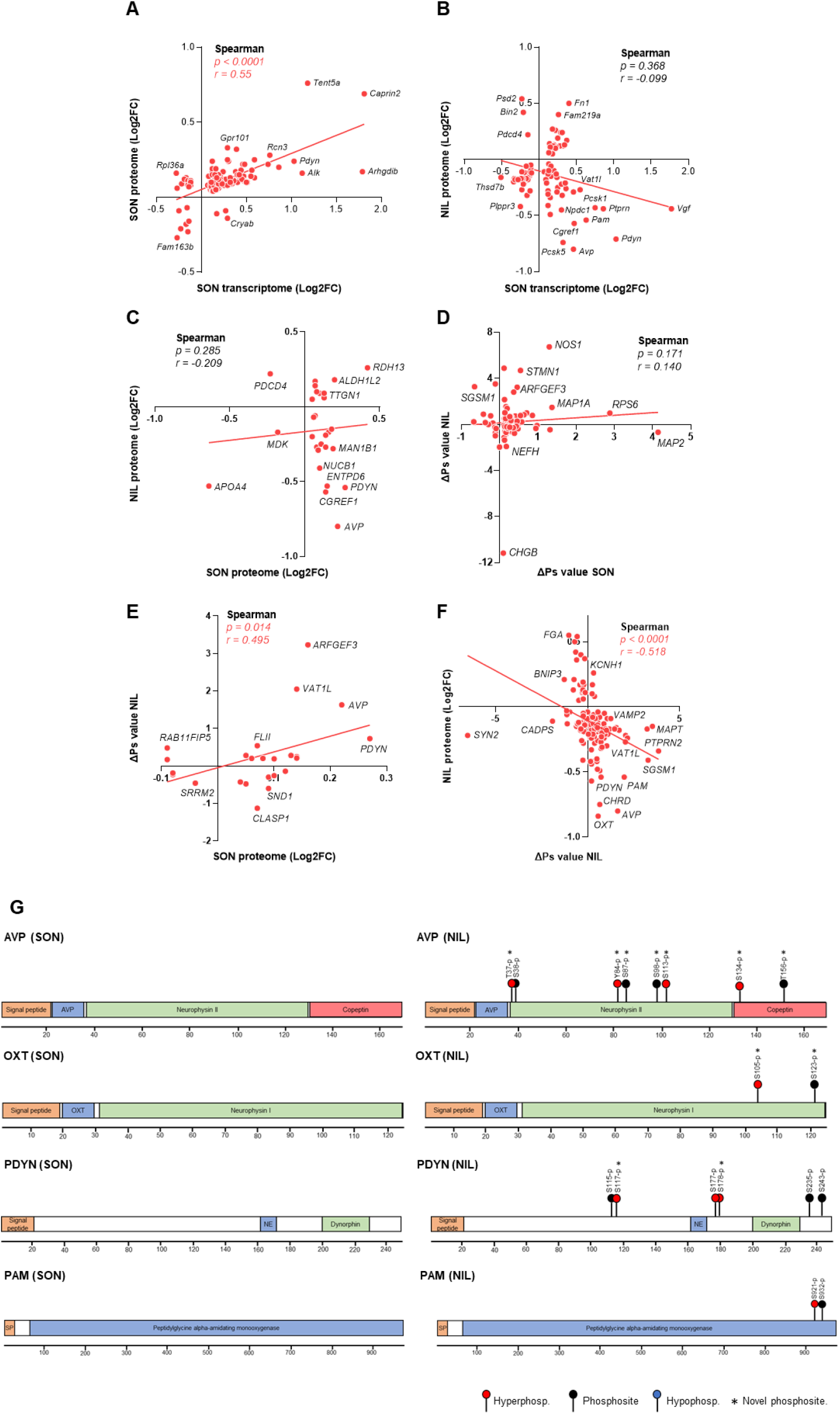
Multi-omic integration of stimulated supraoptic nucleus (SON) and neurointermediate lobe (NIL) (A) Spearman correlation analysis of Log2FC changes in the rat SON proteome and Log2FC changes in the SON transcriptome in response to water deprivation (WD). (B) Spearman correlation analysis of Log2FC changes in the rat NIL proteome and Log2FC changes in the SON transcriptome in response WD. (C) Spearman correlation analysis of Log2FC changes in the rat NIL proteome and SON proteome in response to WD. (D) Spearman correlation analysis of ΔPs changes in the rat NIL and SON in response to WD. (E) Spearman correlation analysis of ΔPs changes in the rat NIL and Log2FC changes in the SON proteome in response to WD. (F) Spearman correlation analysis of Log2FC changes in the rat NIL proteome and ΔPs changes in the NIL in response to WD. (G) Mapping the phosphosites and hyperphosphorylation events in response to WD in Vasopressin-neurophysin 2-copeptin (AVP), Oxytocin-neurophysin 1 (OXT), Proenkephalin-B (PDYN), Peptidylglycine alpha-amidating monooxygenase (PAM) in the SON and NIL.

### Basal state transcriptome vs proteome integration

We then explored the relationship between the transcriptomes and the proteomes of SONs in the basal condition by comparing the proteome data with transcriptomic analysis in Wistar Han rats (Pauža *et al*., 2021). We compared transcripts in the RNAseq dataset with a mean number of reads > 10 with the LC-MS/MS total proteome data from SON and NIL in euhydrated conditions. These comparisons revealed important transcriptome and proteome dynamics in cell bodies and axonal terminals (**Figure 8A, B**). To begin with, there were 6862 transcripts for which the corresponding encoding proteins were not detected by LC-MS/MS. This can be attributed to the lower sensitivity and dynamic range for detection of proteomics and to technical issues such as protein solubility (Dapic et al., 2017). Interestingly, there were 6143 transcripts in the SON which encoded proteins were detected in both the SON and NIL, likely representing a mix of proteins synthesised in the cell body and transported to the axonal terminals and/or synthesised in different cell populations in the SON and NIL. There were 293 proteins exclusively present in the NIL, without their corresponding transcripts in the SON, suggesting either local synthesis of these proteins in the NIL, inputs from non-SON neurones projecting to the NIL, or contamination from blood. There were 1372 transcripts in the SON which encoded proteins present in the NIL but not the SON, possibly reflecting protein transport from cell bodies in SON to axonal terminals in NIL. In addition, there were 852 transcripts with proteins present in the SON, but not the NIL, suggesting that these proteins are synthesised in the SON, but are not transported to the PP, instead having unique biological functions in cells bodies and dendrites, or that they are produced in SON cells other than MCNs.

**Figure 8.**
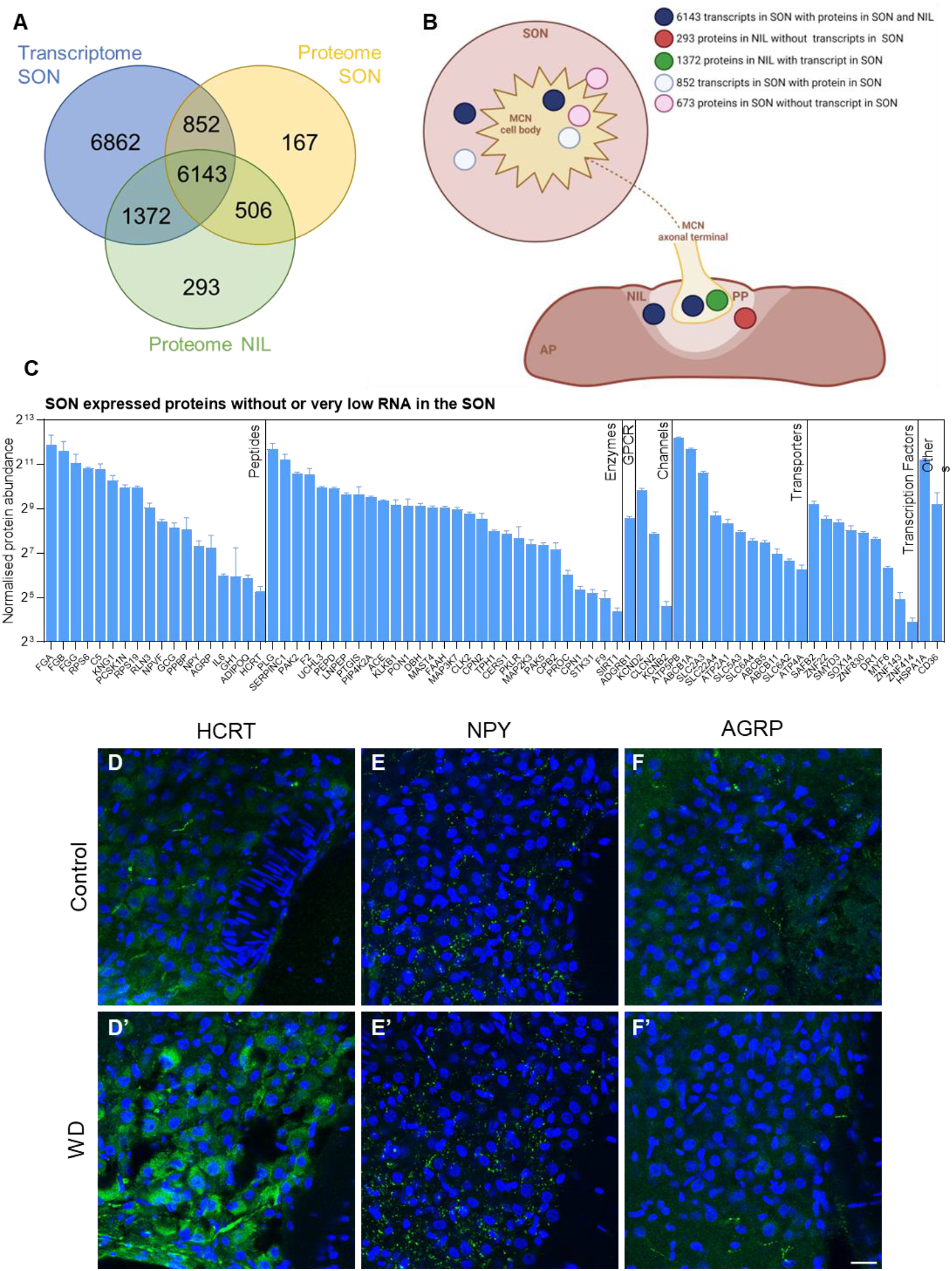
Basal state transcriptome vs proteome integration. (A) Venn diagram showing the number of overlapping proteins in the SON and NIL and genes in the SON in control conditions. (B) Schematic representation of the gene and protein dynamics in the SON and NIL in control conditions according to comparisons from the Venn diagram. Generated using BioRender (https://biorender.com/). (C) Proteins detected in the SON without or very low transcripts in this structure categorised according to their pharmacological classification or their function as a transcription factor. (H) Immunohistochemistry against HCRT, NPY and AGRP in the SON of control and water-deprived (WD) rats. Images are representative of n = 4. Scale bar represents 25 µm.

Interestingly, there were 673 proteins detected in the SON without a corresponding transcript (**Figure 8A, B**). We hypothesised these proteins may provide novel insights regarding SON neuronal circuit connectivity. However, it is important to bear in mind that the discrepancy between RNA and presence of protein could in some cases be due to contamination from proteins present in the blood or to differences in the rat strains used in the transcriptomic and the proteomic analysis. We classified these proteins according to their identity as transcription factors or their physiological or pharmacological categories and included 15 peptides, 23 enzymes, 2 GPCR, 2 channels, 9 transporters and 2 transcription factors (**Figure 8C**). We identified Hypocretin neuropeptide precursor (HCRT) as a candidate peptide being transported into the SON. Interestingly, the protein abundance of HCRT was increased in response to WD. Immunohistochemistry identified HCRT positive afferent fibres (**Figure 8D**), possibly arising from neuronal cell bodies in the lateral and posterior hypothalamus (Date et al., 1999) and was indicative of increased HCRT protein content in the SON (**Figure 8D, D’**). In addition, Neuropeptide Y (NPY), Agouti related protein (AGRP), and Preproglucagon (GCG) were detected at the protein but not mRNA level. The presence of Glucagon-like peptide 1 (GLP1) in afferent fibres within the SON is well-known (Tauchi et al., 2008), possibly arising from the nucleus tractus solitarius (Kabahizi et al., 2022). Immunostaining of NPY revealed axonal terminals containing NPY peptide in the SON (**Figure 8E**) possibly projecting from other hypothalamic structures (Chronwall et al., 1985; Kask et al., 2002). In agreement with LC-MS/MS, no changes in NPY immunolabelling were observed in response to WD (**Figure 8E, E’**). We also detected the presence of ARGP-containing axonal fibres in the SON (**Figure 8F**) and again observed no change in response to WD (**Figure 8F, F’**). Thus, we have identified several signalling proteins that could potentially be transported into the SON from other brain regions to mediate functional outcomes.

## Discussion

In this work we have explored the proteomes and phosphoproteomes of different compartments of the SON and NIL under basal and stimulated conditions. Through comparisons with corresponding SON transcriptomes, we bring a novel perspective to transcriptome, proteome and phosphoproteome dynamics in this uniquely tractable model neuronal system.

Whilst it is well known that proteins synthesised in SON cell bodies are transported to axonal terminals in the PP, our data globally quantifies this phenomenon. When stimulated, MCNs change their steady-state RNA levels and proteomes. In response to WD, proteome and phosphoproteome dynamics differ between the SON and NIL, showing that each neuronal compartment adapts in a very distinctive way to facilitate changes to cell secretory requirements. In the SON, this involves the synthesis of new proteins to meet the demand for newly synthesised peptides (such as AVP and OXT) and associated secretory machinery. The protein phosphorylations in the SON in WD seem to mediate separate functions related to cytoskeleton organisation. In particular, MAP1B S1256 phosphorylation site has already been shown to contribute to the regulation of microtubule dynamics (Trivedi et al., 2005). Also, phosphorylation of the STMN1 phosphosites identified in this work have been shown to promote microtubule stability (Honnappa et al., 2006) and S25-p and S38-p have been found to be phosphorylated in response to hyperosmotic stress (Ng et al., 2010). It has been demonstrated that MNCs have a distinctive cytoskeleton composed of a layer of actin filaments beneath the plasma membrane and a unique network of cytoplasmic actin filaments and microtubule interweaved scaffold (Barad et al., 2020; Prager-Khoutorsky et al., 2014). The cytoskeleton of MCNs undergoes reorganisation in response to hyperosmotic stimuli, which is believed to underlie the intrinsic osmosensitivity of MNCs (Barad *et al*., 2020; Hicks et al., 2020; Prager-Khoutorsky *et al*., 2014). Our data suggests that phosphorylation events in response to WD in the SON may contribute to changes in the cytoskeletal organisation in MCNs. Furthermore, we have also mapped the phosphosites that change in response to WD in proteins involved in cytoskeleton remodelling, such as MAP and other microtubule-organising proteins, providing a global overview of the phosphorylation events mediating cytoskeleton reorganisation.

In the NIL, pathway analysis for changes in the proteome and phosphoproteome in WD informed of adaptations to the synaptic vesicle cycle instrumental for secretion. By mapping phosphosites and responses to stimuli, we provide a global overview of the phosphorylation events involved in the synaptic vesicle cycle and secretion. We report a catalogue of the phosphorylation events that take place in known proteins of the synaptic vesicle cycle, synaptic vesicle clustering, and synaptic exocytosis. These include critical proteins of the synaptic vesicle cycle such as PCLO, BSN and proteins from the VAMP, SNAP, syntaxin, synaptotagmin and synapsin families. Although most of the phosphorylation events that we describe have no currently known functions, some have already been explored. BSN phosphorylation at S2845 has been shown to modulate its anchoring to the presynaptic cytomatrix as part of presynaptic remodelling during synaptic plasticity (Schroder et al., 2013). Phosphorylation of SYN1 at S62/67 causes the dissociation of synaptic vesicles from the actin cytoskeleton resulting on their mobilisation from the reserve pool to the release-ready pool (Chi et al., 2003). Phosphorylation of SNAP25 at T138 inhibits assembly of the SNARE complex and exocytosis (Gao et al., 2016), but increases the size of releasable vesicle pools (Nagy et al., 2004). S14-phosphorylated STX1B has been shown to interact with SNAP25 in specific domains of the axonal plasma membrane with no pools of synaptic vesicles suggesting that these could be fusion sites for a novel class of vesicles beyond traditional active zones. These findings further support the role of these phosphorylation changes in the synaptic vesicle cycle in the HNS. We also provide a catalogue of the phosphorylation events taking place for proteins involved in “presynaptic DCV exocytosis” which include the proteins CADPS, required for Ca^2+^-activated DCV exocytosis (Berwin et al., 1998), RIMS1, DNM1, RAB3A and the syntaxin binding proteins STXBP1 and STXBP5. It has been described that DNM1 is constitutively phosphorylated at S778. This residue is dephosphorylated following neuronal stimulus to facilitate synaptic vesicle endocytosis, which is necessary to maintain a pool of synaptic vesicles within nerve terminals after exocytosis, following which it is rephosphorylated to allow for the next round of synaptic vesicle endocytosis (Tan et al., 2003).

The negative correlation between the ΔPs and the proteome in the NIL as a result of WD suggests that hyperphosphorylation of proteins maybe a key component for neuropeptide processing and/or secretion from nerve terminals. In particular, in the AVP and PDYN precursor proteins we have identified phosphorylation events next to the cleavage sites suggesting that changes in phosphorylation might regulate processing, as it has already been observed for gastrin (Bishop et al., 1998).

The protein NOS1 had high ΔP values both in the SON and the NIL. Interestingly, by mapping the phosphorylation sites and their changes to WD, both in the SON and NIL, we show cell compartment specific phosphorylation patterns (**Figure S2**). We detected hyperphosphorylation events in the flavin mononucleotide (FMN)-binding domain in the SON and NIL as a result of WD, where increased phosphorylation at S847 in the NIL, but not the SON, has been shown to reduce NOS1 activity by inhibiting the binding of Ca^2+^ to the calmodulin domain (Hayashi et al., 1999; Komeima et al., 2000). In addition, NOS1 phosphorylation at S1412 in the NADPH-binding domain (hyperphosphorylated in SON, but not NIL) has been shown to increase the activity of NOS1 (Chen et al., 2021; Khan et al., 2015). It has been suggested that hyperosmotic stimulation induces NO production in MCNs in the SON (da Silva et al., 2013; Reis et al., 2015) reducing AVP and OXT secretion as a feedback compensatory mechanism to prevent over-secretion of these peptides (Pires da Silva et al., 2016). The different phosphorylation events in NOS1 between the NIL and SON identified in this study and the implications regarding the differential activities of this enzyme in discrete cellular structures, can contribute to fully understand the role of NOS1 in MCNs and other neuronal systems.

We have also identified a number of proteins in the SON without the presence of their corresponding transcripts, and validated proteins known to be found in afferents. The SON expresses the HCRT receptors *Hcrtr1* and *Hcrtr2*, the NPY receptors *Npy1r, Npy2r* and *Npy5r*, and the AGRP receptors *Mc3r* and *Mc4r*, the later even increases in response to WD, (Pauža *et al*., 2021) in agreement with regulation by these neuropeptides. The increase in HCRT in WD supports a role for this neuropeptide circuit in the control on MCN functions. Interestingly, HCRT regulates the sleep-wake cycle (Sagi et al., 2021). It has been demonstrated that WD reduces motor activity and increases slow-wave sleep (Martelli et al., 2012), so HCRT could potentially be mediating these effects.

In order to better understand the biological functions of neurones, a comprehensive multiomic understanding of activity-dependent neuronal cellular pathways and processes in distinct cellular and sub-cellular compartments is needed. But in mammals, this is easier said than done. We have taken advantage of the unique anatomical organisation of the HNS to document transcriptome, proteome and phosphoproteome dynamics in this structure in response to neuronal activation (see graphical abstract). These data show how different compartments of the HNS respond to stimulation. This multiomic approach provides a wealth of new knowledge about how neuronal stimulation reshapes the proteome and phosphoproteome to be utilised by the neuroscience community and beyond.

## Materials and Methods

### Animals

All experimental procedures involving animals were performed in strict accordance with the provision of the UK Animals (Scientific Procedures) Act (1986). The study was carried out under a Home Office UK licence (PPL PP9294977) and all the protocols were approved by the University of Bristol Animal Welfare and Ethical Review Board.

Twelve male Sprague-Dawley rats weighing 250–300 g were purchased from Envigo and acclimatised for 10 days. Rats were maintained under a 12:12 light dark cycle (lights on 8.00 am) at a constant temperature of 21-22°C and a relative humidity of 40%-50% with *ad libitum* access to food and water. Rats were housed in groups of 3 with environmental enrichment consisting of nesting material, cardboard tube, and a chew block. Animal cages were randomly assigned to control or WD groups. For the WD group, water was removed for 48 hours with *ad libitum* access remaining for controls.

For proteomic analyses and Western blotting, rats were killed by striking the cranium. The brain was removed from the cranium and placed in an ice-cold brain matrix to separate the forebrain from the hindbrain. The pituitary gland was removed from the base of the cranium and the NIL (containing the PP and the intermediate lobe) dissected from the anterior pituitary. Forebrains and NIL were immediately frozen in powdered dry ice and stored at −80°C. For immunohistochemistry analyses, rats were deeply anesthetised with intraperitoneal administration of sodium pentobarbitone (100 mg/kg,) and transcardially perfused with 0.1M phosphate buffered saline pH 7.4 (PBS) followed by 4% (w/v) paraformaldehyde (PFA) in PBS. The brain and pituitary gland were removed, post-fixed in 4% (w/v) PFA overnight and cryoprotected in 30% (w/v) sucrose in PBS prior to freezing the tissues over liquid nitrogen. All sample collections were performed between 9.00 am and 12.00 pm.

### Protein extraction for proteomic and phosphoproteomic analyses

SON samples were collected bilaterally using a 1-mm micropunch (Fine Scientific Tools) from 100 μm brain coronal sections in a cryostat as described (Greenwood et al., 2014). Proteins from SON and NIL samples were extracted in lysis buffer containing 50 mM Tris-HCl, pH 7.6; 150 mM NaCl; 0.1% (w/v) sodium dodecyl sulfate; 0.5% (w/v) sodium deoxycholate; 1% (v/v) Nonidet P-40; 1 mM EDTA) containing the protease inhibitors 1mM Phenylmethylsulfonyl fluoride (Merck, P7626), Pierce Protease Inhibitor Tablets (Thermo Fisher Scientific, A32963) and Pierce Phosphatase Inhibitor Mini Tablets (Thermo Fisher Scientific, A32957) in three sonication cycles of 12 seconds. Samples were then incubated in ice for 30 minutes, vortexing every 5 minutes, and then centrifuged at 10000 g for 20 minutes at 4°C. The supernatant was transferred to a fresh tube and protein concentrations were determined by the Bradford assay.

### TMT Labelling, High pH reversed-phase chromatography and Phospho-peptide enrichment

Total proteome and phospho proteome analysis were performed at the Bristol Proteomics Facility, University of Bristol. Aliquots of 100 µg of each sample were digested with trypsin (2.5µg trypsin per 100µg protein; 37°C, overnight), labelled with Tandem Mass Tag (TMTpro) sixteen plex reagents according to the manufacturer’s protocol (Thermo Fisher Scientific, Loughborough, LE11 5RG, UK) and the labelled samples pooled.

For the total proteome analysis, an aliquot of 50 µg of the pooled sample was desalted using a SepPak cartridge according to the manufacturer’s instructions (Waters, Milford, Massachusetts, USA). Eluate from the SepPak cartridge was evaporated to dryness and resuspended in buffer A (20 mM ammonium hydroxide, pH 10) prior to fractionation by high pH reversed-phase chromatography using an Ultimate 3000 liquid chromatography system (Thermo Fisher Scientific). In brief, the sample was loaded onto an XBridge BEH C18 Column (130Å, 3.5 µm, 2.1 mm X 150 mm, Waters, UK) in buffer A and peptides eluted with an increasing gradient of buffer B (20 mM Ammonium Hydroxide in acetonitrile, pH 10) from 0-95% (w/v) over 60 minutes. The resulting fractions (20 in total) were evaporated to dryness and resuspended in 1% (v/v) formic acid prior to analysis by nano-LC MSMS using an Orbitrap Fusion Lumos mass spectrometer (Thermo Scientific).

For the Phospho proteome analysis, the remainder of the TMT-labelled pooled sample was desalted using a SepPak cartridge (Waters, Milford, Massachusetts, USA). The eluate from the SepPak cartridge was evaporated to dryness and subjected to TiO2-based phosphopeptide enrichment according to the manufacturer’s instructions (Pierce). The flow-through and washes from the TiO2-based enrichment were then subjected to FeNTA-based phosphopeptide enrichment according to the manufacturer’s instructions (Pierce). The phospho-enriched samples were again evaporated to dryness and then resuspended in 1% formic acid prior to analysis by nano-LC MSMS using an Orbitrap Fusion Lumos mass spectrometer (Thermo Scientific).

### Nano-LC Mass Spectrometry

High pH RP fractions (Total proteome analysis) or the phospho-enriched fractions (Phospho-proteome analysis) were further fractionated using an Ultimate 3000 nano-LC system in line with an Orbitrap Fusion Lumos mass spectrometer (Thermo Scientific). In brief, peptides in 1% (v/v) formic acid were injected onto an Acclaim PepMap C18 nano-trap column (Thermo Scientific). After washing with 0.5% (v/v) acetonitrile 0.1% (v/v) formic acid peptides were resolved on a 250 mm × 75 μm Acclaim PepMap C18 reverse phase analytical column (Thermo Scientific) over a 150 min organic gradient, using 7 gradient segments (1-6% solvent B over 1min., 6-15% B over 58min., 15-32%B over 58min., 32-40%B over 5min., 40-90%B over 1min., held at 90%B for 6min and then reduced to 1%B over 1min.) with a flow rate of 300 nl min^−1^. Solvent A was 0.1% (v/v) formic acid and Solvent B was aqueous 80% (v/v) acetonitrile in 0.1% (v/v) formic acid. Peptides were ionized by nano-electrospray ionization at 2.0kV using a stainless-steel emitter with an internal diameter of 30 μm (Thermo Scientific) and a capillary temperature of 300°C.

All spectra were acquired using an Orbitrap Fusion Lumos mass spectrometer controlled by Xcalibur 3.0 software (Thermo Scientific) and operated in data-dependent acquisition mode using an SPS-MS3 workflow. FTMS1 spectra were collected at a resolution of 120 000, with an automatic gain control (AGC) target of 200 000 and a max injection time of 50ms. Precursors were filtered with an intensity threshold of 5000, according to charge state (to include charge states 2-7) and with monoisotopic peak determination set to Peptide. Previously interrogated precursors were excluded using a dynamic window (60s +/-10ppm). The MS2 precursors were isolated with a quadrupole isolation window of 0.7m/z. ITMS2 spectra were collected with an AGC target of 10 000, max injection time of 70ms and CID collision energy of 35%.

For FTMS3 analysis, the Orbitrap was operated at 50 000 resolution with an AGC target of 50 000 and a max injection time of 105ms. Precursors were fragmented by high energy collision dissociation (HCD) at a normalised collision energy of 60% to ensure maximal TMT reporter ion yield. Synchronous Precursor Selection (SPS) was enabled to include up to 10 MS2 fragment ions in the FTMS3 scan.

### Data processing

The raw data files were processed and quantified using Proteome Discoverer software v2.1 (Thermo Scientific) and searched against the UniProt Rat database (downloaded July 2021: 35859 entries) using the SEQUEST HT algorithm. Peptide precursor mass tolerance was set at 10ppm, and MS/MS tolerance was set at 0.6Da. Search criteria included oxidation of methionine (+15.995Da), acetylation of the protein N-terminus (+42.011Da) and Methionine loss plus acetylation of the protein N-terminus (−89.03Da) as variable modifications and carbamidomethylation of cysteine (+57.0214) and the addition of the TMTpro mass tag (+304.207) to peptide N-termini and lysine as fixed modifications. For the Phospho-proteome analysis, phosphorylation of serine, threonine and tyrosine (+79.966) was also included as a variable modification. Searches were performed with full tryptic digestion and a maximum of 2 missed cleavages were allowed.

### Protein abundance processing

Protein groupings were determined by PD2.2, however, the master protein selection was improved with an in-house script. This enables us to infer biological trends more effectively in the dataset without any loss in the quality of identification or quantification. The MS data were searched against the human Uniprot database retrieved on 2022-01-05 and updated with additional annotation information on 2022-01-20.

The protein abundances were normalised within each sample to total peptide amount, and then scaled to the abundance of the common pool sample (a single representative sample run in each separate TMT experiment) to allow comparisons between experiments. The scaled abundances were then Log2 transformed to bring them closer to a normal distribution.

### Phosphopeptide abundance processing

The phosphorylation status of identified peptide spectral matches (PSMs) was determined by PD2.2 and the site of phosphorylation predicted by PD2.2 using the PhosphoRS module. Phosphorylation sites predicted by PhosphoRS with greater than 70% confidence were taken as the likely phosphorylation site, and phosphopeptides identified with identical sequences and predicted phosphorylation sites, were combined to provide improved quantitation and confidence. The number of PSMs used to calculate the phosphosite abundance is shown in the “Contributing PSMs” column in the excel output.

Where a peptide is predicted to be phosphorylated (based on its mass), but the software is unable to assign the site, the site is listed as “Ambiguous”. Where multiple phosphorylation events are unable to be located to specific sites within a peptide, the word “Ambiguous” is repeated the corresponding number of times.

As peptides can often be matched to multiple proteins, the list of proteins to which each peptide matched was searched against the list of master proteins in the Total Protein analysis, and if a matching protein was identified, this protein was used as the master protein for that peptide.

The experiment was performed by combining an equal amount of each sample, following TMT labelling. 5% of this combined sample was used for the total proteome analysis, and 95% was taken for phosphopeptide enrichment and subsequent analysis of the phosphoproteome. After phospho enrichment we would not necessarily expect the samples to contain the same amount of phosphopeptides, due to biological differences in the levels of phosphorylation between the conditions. As such, normalisation of phosphopeptide abundance was performed using the normalisation factor generated from analysis of the corresponding samples in the total protein dataset. The samples were then scaled to the corresponding pool in the same manner as occurred with the total proteome.

When total protein data is available for the phosphopeptide, the Log2 Scaled Protein abundance was subtracted from the Log2 Scaled Phosphopeptide abundance to adjust the phosphopeptide abundance for any changes in the total protein. As such, if a constitutively phosphorylated protein doubles in its protein abundance as a result of a condition of interest, the Log2 Scaled Phosphopeptide Abundance will show a doubling in the abundance of the phosphopeptide, however the Adjusted Log2 Phosphopeptide Abundance will not show any significant change, as the same proportion of the protein is phosphorylated.

Statistical significance was then determined using Welch’s T-Tests between the conditions of interest. The p-values were false discovery rate (FDR) corrected using the Benjamini-Hochberg method. Data were then exported to excel for ease of use. Since it has been discussed that the use of multiple testing corrected FDR may be too blunt and restrictive for proteomic analysis [18], especially when analysing such an heterogeneous and complex tissue as the brain [19-24], we considered uncorrected p < 0.05 as differentially produced proteins and phosphosites in all our SON and NIL analysis. All data have been deposited at the ProteomeXchange Consortium via the PRIDE partner repository (Perez-Riverol et al., 2019) with the dataset identifier PXD033401.

### Data analysis

Principal component analysis (PCAs) were calculated using the FactoMineR package, and then plotted using the ggplot2 package. Principal Components 1 and 2 were plotted against each other to give an indication of the main sources of variance, and 3 and 4 were plotted to infer any further trends.

In volcano plots, for each comparison the -log10 p-value of each protein was plotted against the log2 fold change using GraphPad Prism 8.4.3. Proteins where p<0.05 are highlighted in blue and red.

Gene Ontology (GO) gene set enrichment analysis was performed using the gProfiler2 package (Kolberg et al., 2020) in R (R core team, 2021, version 4.0.3) (R Core Team, 2021) using a significance threshold of <0.05 Padj for enriched terms. A background list of all detectable proteins produced in each tissue was used. Databases searched included GO:MF, GO:CC, GO:BP and KEGG. Ontologies for the synapse was performed using SynGO (Koopmans *et al*., 2019).

The phosphorylation state change (ΔPs) value calculation was adapted from Wang *et al*. (2018). Briefly, the ΔPs values for protein isoforms encoded by the same gene were determined by the sum of Log2FC of all phosphorylated peptides with statistically significant changes (p < 0.05) in both SON and NIL when comparing the WD and control groups in the phosphoproteome data. We next calculated the average standard deviation (SD) of adjusted log2 normalised abundance from all identified phosphopeptides in the SON (SD = 0.17) and NIL (SD = 0.2). We applied a cut-off at ±0.34 for the SON and ±0.40 for the NIL (>2 SD) to represent the cumulative protein phosphorylation, determining the hyperphosphorylated and hypophosphorylated phosphoproteins on those tissues in response to WD.

We classified changes in the proteome and ΔPs as endogenous peptides, enzymes, G protein-coupled receptors (GPCRs), catalytic receptors, channels, transporters, transcription factors, and other pharmacological targets using the functional classification of the International Union of Basic and Clinical Pharmacology (Harding *et al*., 2022) in association with a manually curated list of validated human transcription factors (Lambert *et al*., 2018). Only the proteins with an existing entry in these databases were catalogued according to this classification. The same approach was used to characterise the newly discovered peptides without, or very low mRNA expressed in the SON.

Mapping of phosphosites and changes in phosphorylation to the protein sequence was done by using PhosphoSitePlus (https://www.phosphosite.org).

The relationship between basal and stimulated transcriptomes and proteomes was examined by comparing the proteome output obtained in the present study in basal conditions and after 48 h WD with previous SON transcriptomic analysis in Wistar Han rats in basal conditions and following 72 h WD (Pauža et al., 2021). It is important to note that SONs were collected by the same researcher minimising variability during sample collection and processing. To rule out the impact of strain, we compared Wistar Han rats (Pauža et al., 2021) and Sprague-Dawley rats RNAseq datasets (unpublished) and found that SON transcriptomes from both rat strains were highly correlated (Spearman r = 0.940, p-value < 0.0001).

### Immunohistochemistry

Coronal sections of the forebrain containing the hypothalamus and NILs were cut on a cryostat at 40 μm and kept in PBS at 4°C. To prevent nonspecific protein binding, sections were blocked in PBS containing 0.3% (v/v) Triton X-100, 4% (w/v) bovine serum albumin (BSA), and 5% (v/v) donkey serum at RT for 1 hour. Following this, the NIL sections were incubated overnight at 4°C with the primary antibodies against NOS1 (1:200, Santa Cruz biotechnology, sc-5302), phospho-nNOS (Ser852) (1:100, Thermo Fisher Scientific, PA5-38305), phospho-Synapsin I (Ser62, Ser67) (1:200, Millipore, AB9848), phospho-Synapsin 2 (Ser425) (1:200, Thermo Fisher Scientific, PA5-64855), and Synapsin (1:200, Cell Signaling Technology, 2312S) in PBS containing 0.3% (v/v) Triton X-100, 4% (w/v) BSA, and 1% (v/v) donkey serum. Tissue sections containing the SON were incubated with antibodies against NOS1 (1:200, Santa Cruz biotechnology, sc-5302), Orexin A/Hypocretin-1 (1:1000, R and D Systems, MAB763) Phospho-JUND (Ser255) (1:100, Thermo Fisher Scientific, PA5-104821), Phospho-nNOS (Ser1417) (1:500, Thermo Fisher Scientific, PA1-032), Phospho-Stathmin 1 (Ser24) (1:200, MyBioSource, MBS9600965), Phospho-S6 Ribosomal Protein (Ser240/244) (1:500, Cell Signaling Technology, 5364S), Stathmin 1 (1:500, GeneTex, GTX104707) and S6 Ribosomal Protein (1:100, Cell Signaling Technology 2317S) in PBS containing 0.3% (v/v) Triton X-100, 4% (w/v) BSA, and 1% (v/v) donkey serum. All antibodies were incubated at 4°C overnight, with the exemption of the antibodies against Orexin A/Hypocretin-1 and Phospho-Stathmin 1 (Ser24) that were incubated for 48 hours. Following incubation with the primary antibodies, sections were washed in PBS and incubated with the secondary antibodies donkey anti-rabbit Alexa Fluor Plus 488 (Thermo Fisher Scientific, A32790) and donkey anti-mouse Alexa Fluor 594 (Molecular Probes, A-21203) at a 1:500 dilution in PBS containing 0.1% (v/v) Triton, 4% (w/v) BSA, and 1% (v/v) donkey serum at RT for 1 h. Then, the sections were washed in PBS, incubated with 4’,6-Diamidino-2-phenylindole dihydrochloride (DAPI, D1306; Molecular Probes) in PBS and mounted with ProLong Gold Antifade Mountant (Thermo Fisher Scientific, P36930).

Images were acquired using a Leica SP5-II AOBS confocal laser scanning microscope attached to a Leica DMI 6000 inverted epifluorescence microscope with a 20x and a 63x PL APO CS lens. Raw image files were processed to generate composite images using the open access image analysis software, Fiji.

### Western Blot

Protein extraction from the NIL was performed as described (Greenwood et al., 2015a). Protein samples were prepared to 1× Laemmli buffer solution (2% w/v SDS, 10% v/v glycerol, 5% v/v 2-mercaptoethanol, 0.002% w/v bromophenol blue and 0.125 M Tris HCl, pH 6.8). Samples were heated at 95°C in a hot block for 5 minutes. For semiquantitative analysis of protein levels, 20 μg/lane of total protein (determined in duplicate by Bio-Rad Protein Assay with BSA as standards) was loaded for control and WD samples. Proteins were fractionated on 8% (w/v) sodium dodecyl sulfate polyacrylamide gels and transferred to Immobilon^®^-P PVDF Membrane (Merck, IPVH00010). Membranes were incubated in 5% (w/v) BSA in Tris-buffered saline (150 mM NaCl; 20 mM Tris-HCl, pH 7.6) with 0.1% (v/v) Tween 20 (TBS-T) for 1 hour followed by incubation with the primary antibodies NOS1 (1:200, Santa Cruz biotechnology, sc-5302), phospho-nNOS (Ser852) (1:1000, Invitrogen, PA5-38305), phospho-Synapsin I (Ser62,Ser67) (1:1000, Merk, AB9848), phospho-Synapsin 2 (Ser425) (1:5000, Invitrogen, PA5-64855), Synapsin (1:1000, Cell Signaling Technology, 2312S) and Tubulin (1:5000, Covance, MMS-489P) overnight at 4°C. Following three washes in TBS-T, the membranes were incubated with the appropriate secondary antibody conjugated with horseradish peroxidase for 1 hour. After three washes in TBS-T, the signal was detected using chemiluminescence SuperSignal West Dura Extended Duration Substrate reagent (Thermo Fisher Scientific, 34075). Immunoblots were stripped in Restore Western Blot Stripping Buffer (Thermo Fisher Scientific, 21059) and reprobed to assess the multiple proteins in the same blot.

### Statistical analysis

Statistical analyses were performed with GraphPad 8.4.3 Software. For western blot signal quantification, assessment of the normality of data was performed by Shapiro-Wilk test. Means between two groups were compared using independent-sample unpaired Student’s t tests where data are expressed as box and whisker plots. For the P-SYN1 and P-SYN quantifications, one sample was excluded from the analysis due to the absence of total SYN signal (**Figure S1**), otherwise all samples were included in the analysis. Spearman correlation analysis were also performed in GraphPad Prism Prism 8.4.3. In all cases p < 0.05 was considered statistically significant.

## Data and code availability

All data are available in the manuscript or supplemental information. The LC-MS/MS proteomics and phosphoproteomics data have been deposited to the ProteomeXchange Consortium via the PRIDE (Perez-Riverol *et al*., 2019) partner repository with the dataset identifier PXD033401. Any additional information is available from the lead contact upon request.

## Acknowledgments

The authors gratefully acknowledge Dr Kate J. Heesom and Dr Philip A Lewis from the University of Bristol Proteomics Facility for their support and assistance. We also thank the Wolfson Bioimaging Facility. This research was supported by grants from the Biotechnology and Biological Sciences Research Council (BBSRC; BB/R016879/1) to D.M., S.B.L. and M.P.G. and from the Leverhulme Trust (RPG-2017-287) to D.M. and M.P.G. Students were supported by grants from the Biotechnology and Biological Sciences Research Council-SWBio DTP programme (BBSRC BB/M009122/1) to B.T.G., and the British Heart Foundation to A.G.P (BHF FS/17/60/33474) and to N.B. (FS/4yPhD/F/21/34162).

---

**Supplementary Figure 1.**
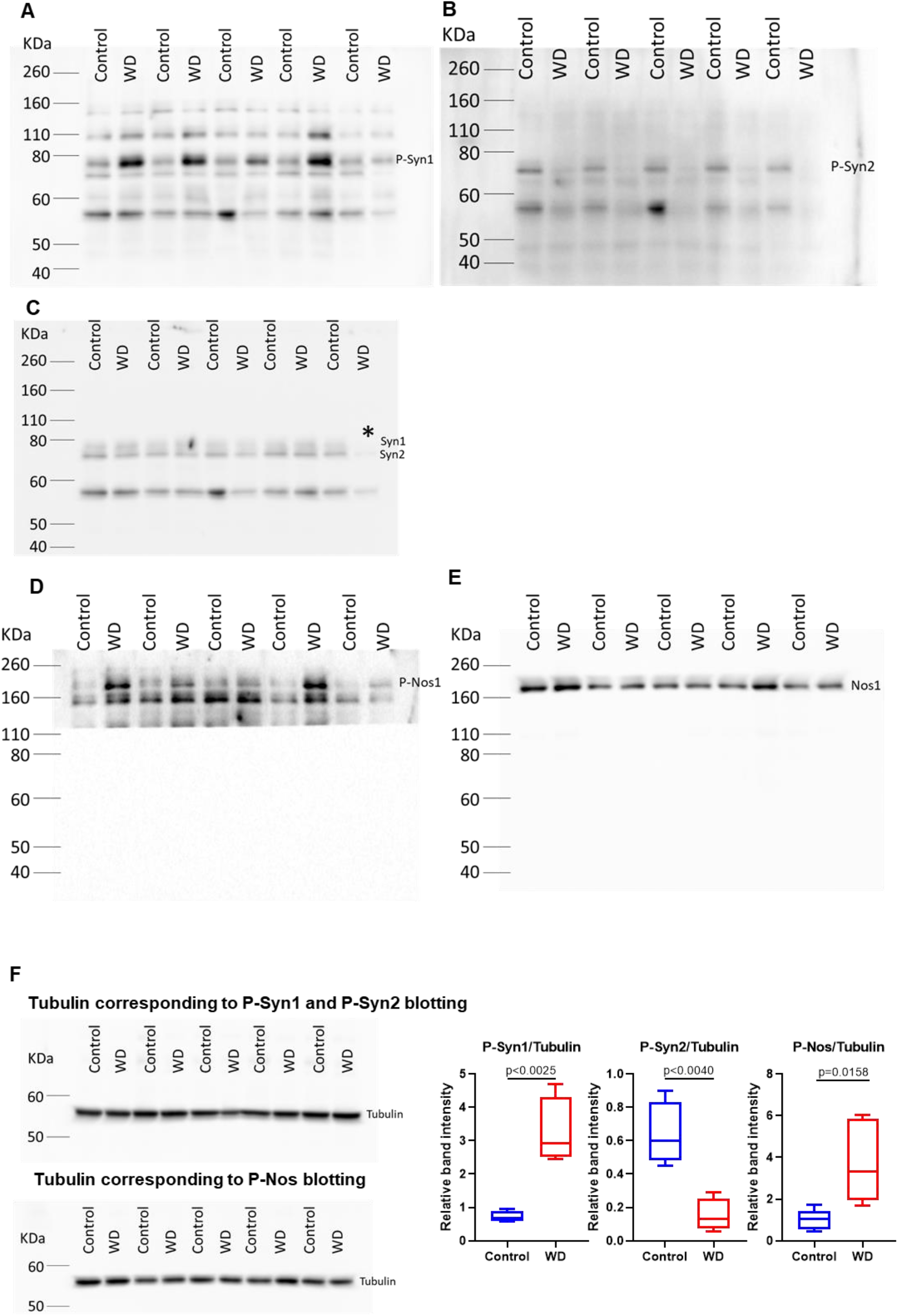
Complete Western Blots with control and 48 hours water deprived (WD) blotted against P-Syn1 (A), P-Syn2 (B), Syn (C), P-Nos (D) and Nos (E). (C) Due to the lack of Syn signal in the last lane (*), this sample was excluded from the P-Syn1 and P-Syn2 analysis. (D) This membrane was covered during signal detection to prevent signal bleed-through from previous rounds of protein detection. (F) Normalising the signal to tubulin rendered the same results as normalising to total Syn and total Nos.

**Supplementary Figure 2.**
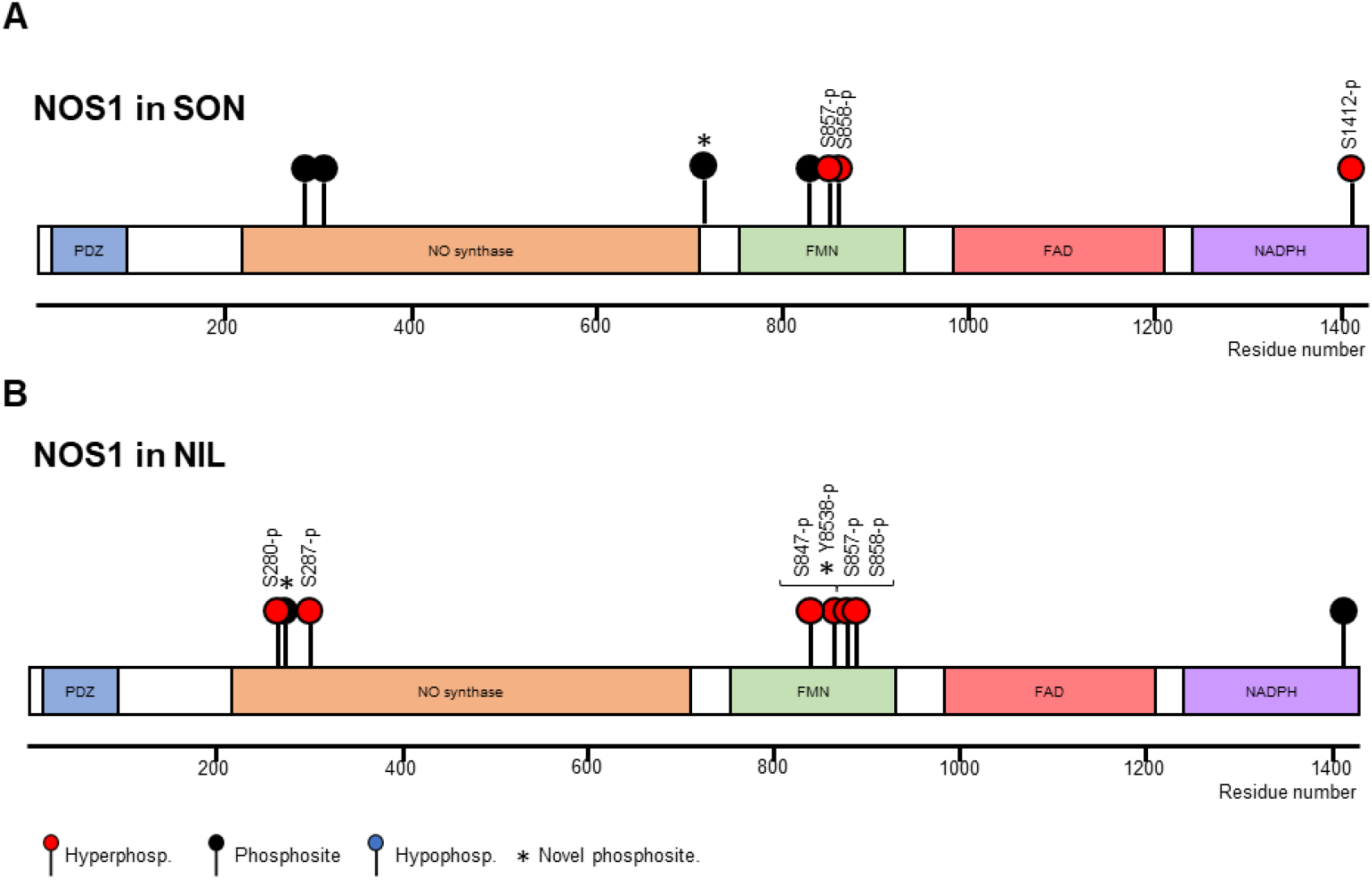
Mapping the phosphosites and hyper and hypophosphorylation events in NOS1 in response to water deprivation in the (A) SON and (B) NIL.

